# A Flat Protein Complex Shapes Rough ER Membrane Sheets

**DOI:** 10.1101/2023.10.06.559866

**Authors:** Eric M. Sawyer, Liv E. Jensen, Janet B. Meehl, Kevin P. Larsen, Daniel A. Petito, James H. Hurley, Gia K. Voeltz

## Abstract

Rough ER sheets are a fundamental domain of the ER and the gateway into the secretory pathway. While reticulon proteins stabilize high-curvature ER tubules, it is unclear if other proteins scaffold the flat membranes of rough ER sheets. Through a proteomics screen using ER sheet localized RNA-binding proteins as bait, we identify the Sigma-1 receptor (SigmaR1) as an ER sheet shaping factor. High-resolution live cell imaging and electron tomography assign SigmaR1 as an ER sheet-localized factor whose levels determine the amount of rough ER sheets in cells. Structure-guided mutagenesis and in vitro reconstitution on giant unilamellar vesicles further support a mechanism whereby SigmaR1 oligomers use their extended arrays of amphipathic helices to bind and flatten the lumenal leaflet of ER membranes. Our results demonstrate an unexpected way for proteins to sense and propagate flat membrane sheets.

## INTRODUCTION

The endoplasmic reticulum (ER) exists as a single, continuous membrane network divided into specialized and morphologically distinct domains. Besides the nuclear envelope, the peripheral ER is divided into two main structural and functional domains: smooth ER tubules and rough ER sheets.^1^ Smooth ER tubules function to regulate lipid synthesis, ion transport, and various functions at contact sites with other organelles,^2^ whereas rough ER sheets provide the location of translation and translocation of membrane and secreted proteins.^1^ The functional balance between these two domains is clear from electron microscopy studies.^3,4^ Professional secretory cells reveal dramatic expansion of layers of rough ER sheets.^5^ Conversely, smooth ER tubules predominate in cells specialized for steroid hormone biosynthesis.^5^ However, many eukaryotic cells have an abundance of both domains that are located within a single expansive membrane network.^1,4–6^ How a membrane bilayer can be molded into contiguous and yet structurally distinct domains that are specialized for local functions is a mechanistic challenge.

Factors have been identified that regulate the biogenesis of ER tubules, for example the reticulon family of ER proteins found in all eukaryotic cells.^7^ Reticulons and reticulon-like proteins contain conserved hairpin membrane domains, which wedge into the cytosolic leaflet of the ER membrane to induce the membrane curvature required for tubule stability.^7–10^ Microtubule (MT)-dependent processes also produce dynamic ER tubules in animal cells. At least two different MT-dependent mechanisms exist for ER tubule dynamics: “sliding,” a process whereby ER tubules move along stable MTs using molecular motors, and tip attachment complex (TAC), whereby ER tubules are linked to microtubule-associated proteins (MAPs) and grow and shrink in association with dynamic MT tips.^11,12^

By comparison, much less is understood about mechanisms that flatten ER membranes into rough ER sheets. Reticulons stabilize the curved edges of sheets, and altering the ratio of lipids to reticulons is sufficient to shift the balance between sheets and tubules.^3,7,13–15^ The most studied ER protein linked to ER sheet biogenesis is Climp63. However, Climp63 is localized to both tubules and sheets, and data suggest that it regulates the structure of both by forming a linker/tether that traverses the ER lumen to define the width of sheets and wider tubules.^15–18^ The cytoplasmic domain of Climp63 has a binding site for microtubules, but it is not clear how these two functions are coordinated.^19,20^ Indeed, overexpression of Climp63 reliably drives the formation of more sheets.^15,17,21,22^ However, its depletion does not reduce sheet abundance, though it does narrow sheet diameter, which is why it is considered a lumenal spacer.^15^ Thus, there is currently no ER membrane protein known to directly stabilize flat membrane bilayers to oppose the reticulons’ ability to stabilize membrane curvature. Here we used candidate-based and proteomics approaches to identify a key factor that directly shapes rough ER sheets by a unique mechanism that functions from the lumenal surface of the ER membrane to oppose membrane curvature and tubule formation.

## RESULTS

### ER RNA-binding proteins reside in and expand ER sheets

Our goal was to identify a rough ER sheet-shaping protein. Our strategy assumed that such a protein might be closely associated with mRNA and ribosomes since each are defining functional features of the rough ER. Among a handful of ER integral membrane RNA-binding proteins^15,23–28^ are two proteins with poorly understood molecular functions, Lyric/Mtdh/Aeg-1 and Lrrc59/p34. Lyric was previously shown to interact preferentially with mRNAs encoding membrane/secreted proteins.^23,28^ Lrrc59 (previously called p34) has ribosome-binding activity in a reconstituted system.^27,29^ Proteomic and transcriptomic studies are consistent with involvement of Lrrc59 in an ER translational process, possibly via the signal recognition particle/SRP.^24,30^ We speculated that these proteins might localize to or function to organize rough ER sheet domains, which are enriched with translated mRNAs. First, as a control, we co-transfected the general ER marker BFP-KDEL with mNeon-Sec61β (Figure 1A), which is already known to mark all ER domains when overexpressed.^8^ We scored ER sheet levels by directly measuring ER sheet area using an automated image analysis pipeline that distinguishes ER sheets from tubules by their shapes (Figure 1B, Figure S1A-E). Then, we applied the same analysis to cells transfected with mNeon-tagged Lyric or Lrrc59 over a titration series of expression levels (Figure 1D-F). Lyric and Lrrc59 were both enriched in ER sheets across expression levels and expanded ER sheets at higher expression levels (Figure 1B, D-E). Sheet enrichment persisted even when highly overexpressed (Figure 1D-E). A shared characteristic of both proteins is an extended cytosolic domain enriched with basic amino acids (Figure 1C), which might mediate nucleic acid binding.^23^ Our results demonstrate that Lyric and Lrrc59 are sheet-specific ER RNA-binding proteins that promote ER sheet expansion.

**Figure 1.**
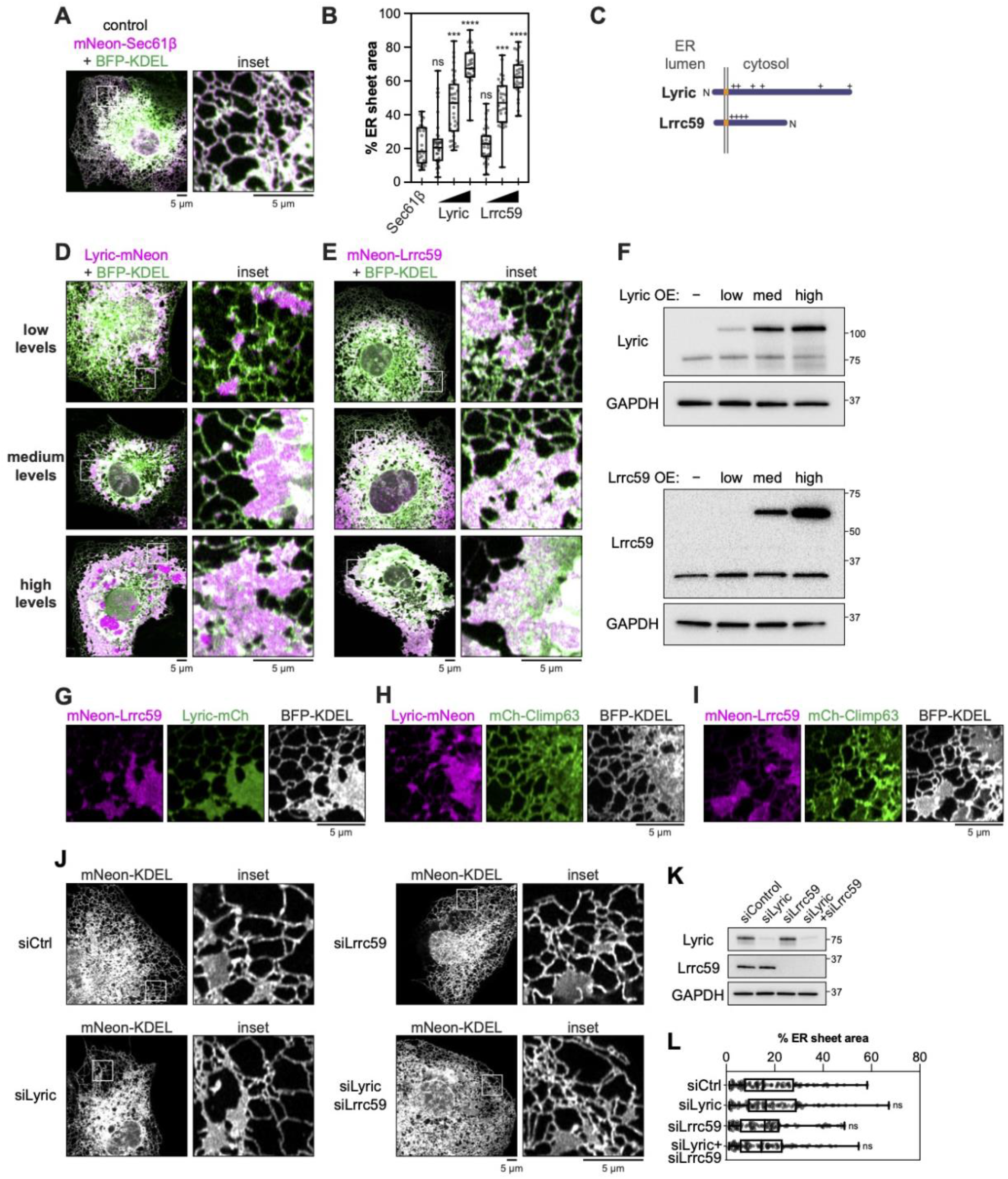
ER RNA-binding proteins Lyric and Lrrc59 reside in and expand ER sheets. (A) Representative merged images of Cos-7 cells co-transfected with a lumenal ER marker BFP-KDEL (green) and mNeon-Sec61β (control, magenta). (B) Quantification of ER sheet area expansion due to ER RBP expression—protocol described in Fig S1. *n* ≥ 30 cells per treatment pooled from 3 biological replicates. Kruskal-Wallis test with Dunn’s multiple comparisons test of each condition compared to the Sec61β control: **** p < 0.0001, *** p < 0.001, ns p ≥ 0.05 (C) Cartoon depicting the topology of Lyric and Lrrc59. +, high density of basic amino acids. (D-E) Representative merged images of Cos-7 cells co-transfected with BFP-KDEL and mNeon-tagged Lyric or Lrrc59. (F) Immunoblot analysis of Cos-7 cells transfected with low, medium, or high plasmid amounts matching the conditions in (D-E). (G-I) Images of Cos-7 cells co-transfected with BFP-KDEL (white) and combinations of Climp63, Lyric, and Lrrc59. (J) Representative images of ER morphology in Cos-7 cells co-transfected with mNeon-KDEL and siRNAs to deplete Lyric, Lrrc59, or both, relative to an siRNA control. (K) Immunoblot analysis of depletion of Lyric or Lrrc59 by siRNA treatment in experiment from (J). (L) Quantification of ER sheet area in control and depletion, as described previously. *n* ≥ 95 cells per treatment from 3 biological replicates. Kruskal-Wallis test with Dunn’s multiple comparisons test of each condition to the siRNA control: ns, p ≥ 0.05.

Next, we co-transfected Cos-7 cells with low levels of Lyric, Lrrc59, and BFP-KDEL to test if they co-localized to the same domains. Lyric and Lrrc59 localized to the same ER sheets when simultaneously expressed (Figure 1G). Interestingly, neither Lyric nor Lrrc59 co-localized with the Climp63 lumenal spacer protein either in ER tubules or sheets (Figure 1H-I).^15^ Our data reveal that Lyric and Lrrc59 have features that would be expected for a protein that contributes to the biogenesis of ER sheets: they localize exclusively to ER sheets at all expression levels and at higher levels expand ER sheets and reduce ER tubules (Fig 1B, D-F).

Next, we tested whether Lyric and/or Lrrc59 depletion would alter the abundance of ER sheets in cells. We co-transfected Cos-7 cells with a general ER marker to visualize ER morphology (mNeon-KDEL) and with either control, Lyric, Lrrc59, or Lyric and Lrrc59 siRNAs (Figure 1J). ER sheet abundance/area was not altered by depletion of either Lyric or Lrrc59 or even by the combined Lyric/Lrrc59 depletion (Figure 1J and 1L). Immunoblot staining confirmed that protein depletion was highly efficient (Figure 1K). Thus, although Lyric and Lrrc59 are ER sheet localized proteins whose expression can drive ER sheet expansion, they are not required to maintain normal levels of ER sheets in cells.

### Lrrc59 binding partners include ribosomal proteins and rough ER proteins

We reasoned that since Lyric and Lrrc59 localize exclusively to ER sheets, can promote ER sheet expansion, and have been linked to RNA/ribosome binding,^23,24,27–30^ they might be useful handles to co-purify a factor required to maintain normal levels of rough ER sheets in cells. We generated HeLa cell lines stably expressing 3x FLAG-mNeon-tagged Lyric or Lrrc59 to co-purify rough ER sheet proteins, compared to an untagged control (Figure 2A). We subjected these cell lines to gentle lysis with 1% digitonin and FLAG immunoprecipitation using a protocol known to recover intact ER membrane protein complexes (Figure 2A). First, we confirmed efficient recovery of tagged Lyric and Lrrc59 bait proteins by immunoprecipitation (see immunoblot analysis, Figure 2B). Colloidal Coomassie stain of co-immunoprecipitated proteins showed unique banding profiles (Figure 2B). Co-immunoprecipitation samples were subsequently analyzed by LC-MS/MS. We used the iBAQ label-free quantification method to estimate the abundance of proteins in the control, Lyric, and Lrrc59 IP samples (Table S1). We imposed a 10-fold cutoff in bait/untagged control iBAQ score ratios to filter out background and plotted the remaining hits based on their absolute iBAQ scores from the Lyric or Lrrc59 IP, ordered by rank (Figure 2C-D). The majority of the hits in both cases were ribosomal proteins (plotted with lavender circles). Among the other top hits were several proteins with known rough ER functions: SEC61B (Sec61β) is a component of the Sec61 translocon, SSR4 (TRAPδ) is a component of the translocon-associated TRAP complex, and SRPRB (SRβ) is a component of the SRP receptor. As ribosomes are the defining characteristic of “rough” ER, and other top hits have well-established rough ER functions, we judged our data to be reliable.

**Figure 2.**
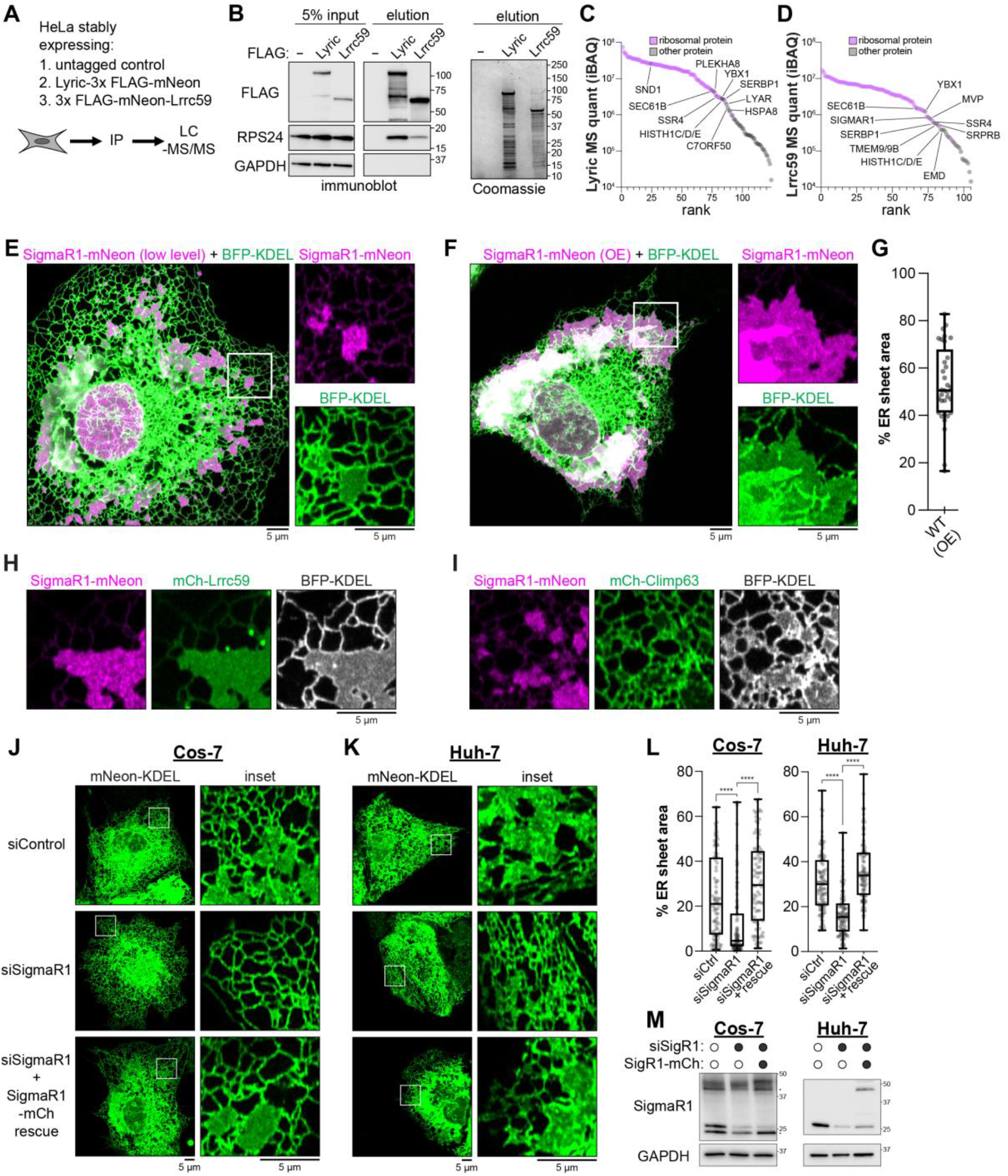
SigmaR1 is an ER sheet-specific protein. (A) Schematic of immunoprecipitation experiment. (B) Analysis of input and elution fractions by immunoblot and colloidal Coomassie-G250 stain. (C) Label-free mass spectrometry quantifications (iBAQ) of Lyric co-precipitated proteins at least 10-fold enriched relative to the untagged control. (D) Lrrc59 co-precipitated proteins analyzed identical to (C). (E-F) Representative merged images of Cos-7 cells transfected with an ER marker (BFP-KDEL) and low (E) or higher levels (F) of SigmaR1-mNeon. (G) Quantification of ER sheet area due to SigmaR1 overexpression in (F). (H-I) Representative merged images of Cos-7 cells co-transfected with BFP-KDEL and low levels of SigmaR1-mNeon with low levels of mCh-Lrrc59 (H) or mCh-Climp63 (I). (J-K) Representative images of Cos-7 cells (J) or Huh-7 cells (K) co-transfected with mNeon-KDEL and indicated combinations of control or SigmaR1 siRNAs and SigmaR1-mCh rescue plasmid. (L) Quantification of ER sheet area in Cos-7 and Huh-7 cells from (J-K). *n* ≥ 89 cells per treatment pooled from 3 experiments. Kruskal-Wallis test with Dunn’s multiple comparisons test: **** p < 0.0001. (M) Immunoblot analyses (anti-SigmaR1) of lysates from (J-K). Representative of 3 biological replicates. * band of unknown identity.

### Sigma-1 receptor levels determine the amount of ER sheets

One of the top hits in the Lrrc59 IP was SIGMAR1 (Sigma-1 receptor, or SigmaR1). SigmaR1 is an integral ER membrane protein first identified by its ability to bind opioid drugs— an ability that was later refuted.^31^ SigmaR1 has also been implicated in controlling ER calcium signaling functions through store-operated calcium entry and IP3 receptors.^31–33^ It has also been proposed to act as a chaperone based on co-immunoprecipitation with putative clients.^32^ This function seems unlikely because SigmaR1 does not contain any conventional chaperone or ATPase domains. SigmaR1 also shares sequence homology with the fungal sterol isomerase, Erg2, though this as a function that has also been refuted, as SigmaR1 is catalytically inactive.^31^ In humans, the same enzymatic step has been co-opted by the structurally unrelated enzyme EBP.^34–36^ The numerous pleotropic effects associated with SigmaR1 could, similar to reticulons, all be the outcome of altering ER shape. However, most compelling is the molecular structure of SigmaR1, which has been solved.^37^ The structure^37^ reveals a homotrimeric complex that resides in the ER lumen^38^ and contains multiple structural features that we will discuss because these features provide insight into its proposed mechanism for flattening membranes.

First, we determined SigmaR1 localization by co-transfecting Cos-7 cells with SigmaR1-mNeon along with an ER marker (BFP-KDEL). SigmaR1 localized to ER sheets, and partitioned away from ER tubules, at both low and higher expression levels (Figure 2E and 2F). Similar to the Lrrc59 bait, SigmaR1 expression also led to the expansion of ER sheets, and a reduction in ER tubules (Figure 2F-G). SigmaR1 also co-localized with the Lrcc59 bait protein in ER sheets and partitioned away from tubules at low and high levels (Figure 2E-H). Like Lrrc59, SigmaR1 partitioned away from both tubule and sheet domains containing the Climp63 lumenal linker protein (Figure 2I). Thus, like Lrcc59 (and Lyric), SigmaR1 is a bona-fide ER sheet protein, and its expression leads to further ER sheet expansion.

Next, we asked if SigmaR1 depletion reduced the amount of ER sheets in cells. SigmaR1 was efficiently depleted by siRNAs in Cos-7 cells, and we scored ER morphology by co-transfection with mNeon-KDEL (Figure 2J, L-M). SigmaR1 depletion significantly reduced the amount of ER sheets in cells relative to a control siRNA, and ER sheets were restored by re-expression of an siRNA-resistant SigmaR1-mCherry transgene (Figure 2J, L-M, Figure S2A). To confirm the robustness of this finding, we replicated these experiments in Huh-7 cells. Huh-7 cells are hepatocyte-derived and have a naturally high proportion of ER sheets, presumably due to their secretory cell functions. SigmaR1 depletion from Huh-7 cells led to a striking reduction in ER sheet area that was fully and quantitatively rescued by re-expression of the siRNA-resistant SigmaR1-mCherry construct (Figure 2K-M, Figure S2B). We conclude that SigmaR1 is required for ER sheet morphology in cells.

### SigmaR1 promotes the biogenesis of rough ER sheets

Although classic EM studies clearly defined rough ER sheets as extended, flat membrane domains,^5^ recent studies have identified additional ER shapes characterized by fenestrations or high densities of clustered ER tubules.^39,40^ To clarify whether SigmaR1-marked ER sheets are bona fide sheets, we analyzed them at nm-scale resolution by electron microscopy and tomography of high-pressure frozen cells. We generated Cos-7 cell lines stably overexpressing mNeon-tagged SigmaR1 or an empty vector control and processed each cell line by high pressure freezing and freeze substitution to preserve membrane ultrastructure for analysis by electron tomography. SigmaR1 overexpressing cells contained a dramatic expansion of ER sheets, as seen in tomographic slices (Figure 3A, Movies S1-S2). SigmaR1-induced ER sheets were ribosome-studded, unfenestrated, and extended uninterrupted through large cellular volumes (Figure 3A, Movies S1-S2). We measured the lengths of rough ER membrane profiles across a series of sections spanning the reconstructed 3D volumes and found that SigmaR1 overexpression led to a significant expansion of rough ER membranes (Figure 3B). Even though the amount of ER was expanded, the ER membranes remained densely ribosome studded (Figure 3A). Our results demonstrate that SigmaR1 drives proliferation of classical rough ER sheets.

**Figure 3.**
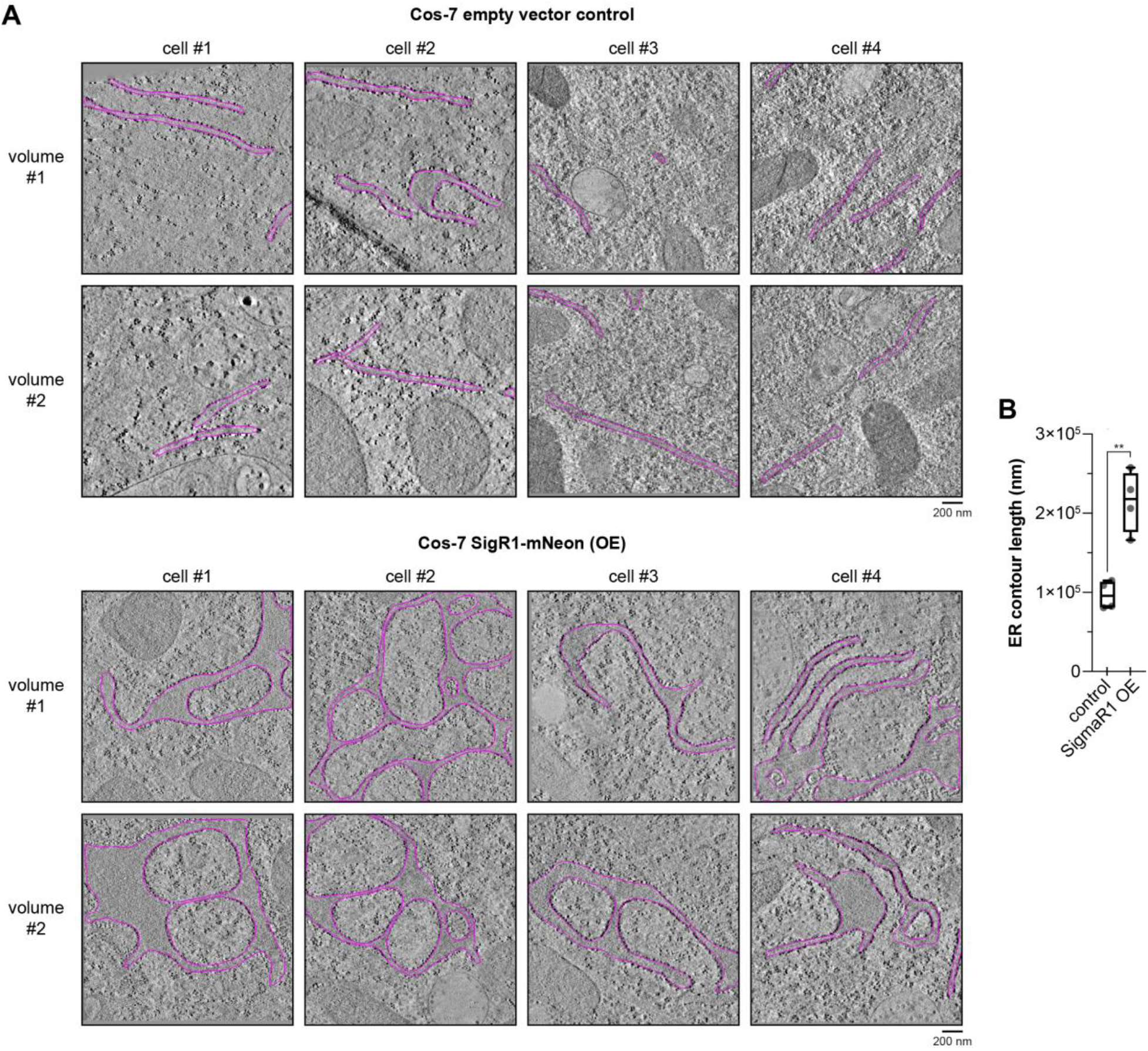
SigmaR1 is sufficient to assemble extended rough ER sheet domains. (A) Representative tomographic slices of high-pressure frozen and freeze substituted Cos-7 cells expressing an empty vector control or SigmaR1-mNeon. Tomographic slices from two different XY positions are shown for each cell with ER membrane contours in magenta. (B) Quantification of ER membrane contour lengths from the experiments in (A)—see Materials and Methods. *n* = 4 cells per treatment from 2 biological replicates. Unpaired t-test ** p < 0.01.

### The SigmaR1 lumenal trimer is sufficient to expand ER sheets

Previous x-ray crystallography studies solved the structure of SigmaR1 as a homotrimer (Figure 4A).^37^ Each subunit is a ∼25 kD single-pass integral membrane protein that terminates with two alpha helices at its C terminus (Figure 4B). These alpha helices have amphipathic character and expose hydrophobic amino acids to the membrane-facing surface of the complex (Figure 4B). Multiple experimental approaches confirm that SigmaR1 is oriented as an N_cyt_ (or type II) topology membrane protein.^38^ This orientation positions the large majority of SigmaR1 within the ER lumen, where its amphipathic helices could lie flat against the inner leaflet of the membrane (Figure 4A-C).

**Figure 4.**
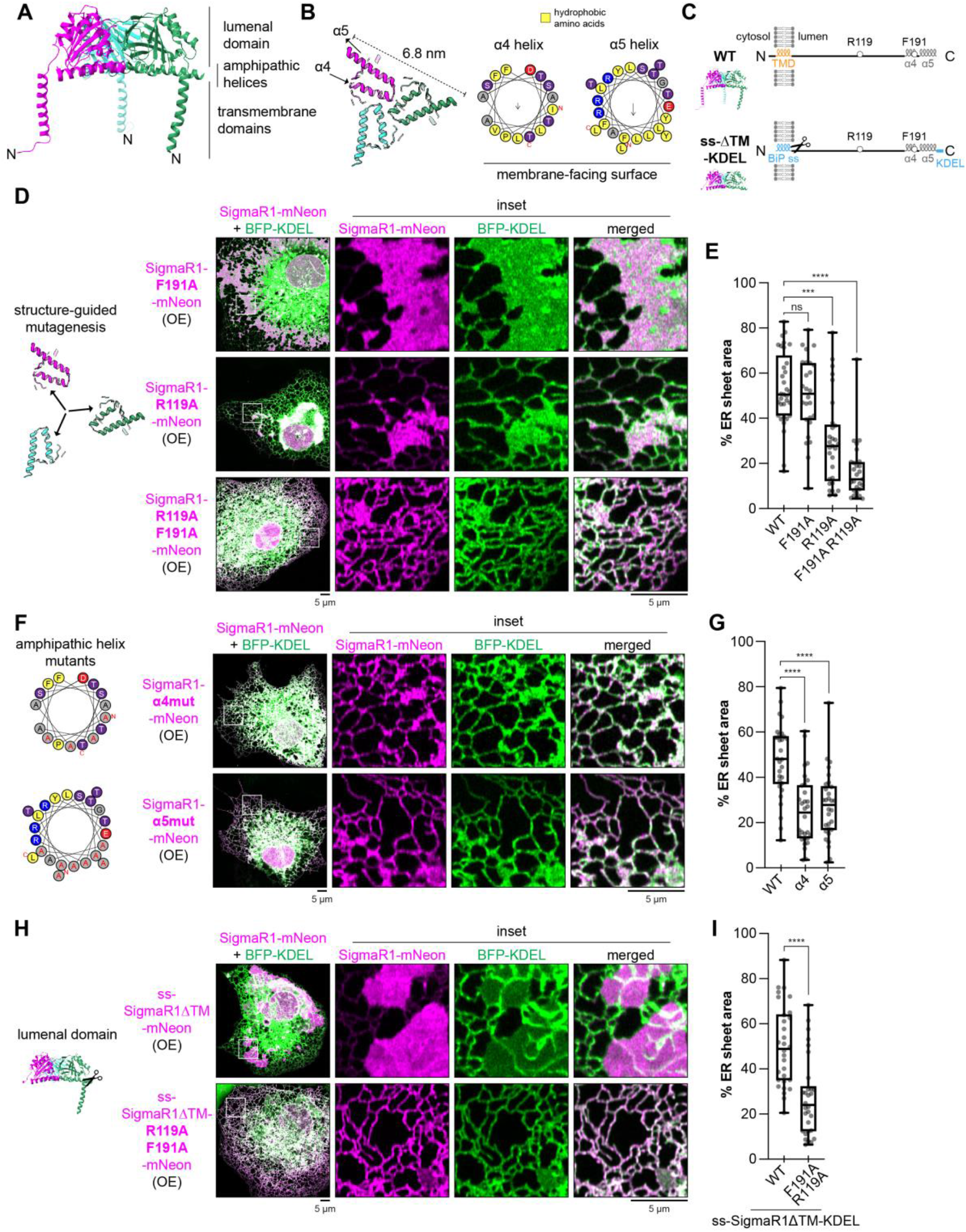
Assembly of the SigmaR1 lumenal trimer drives ER sheet biogenesis. (A) Published^37^ crystal structure (PDB:5hk1) of SigmaR1 trimer colored by polypeptide chain. (B) Membrane-facing surface of SigmaR1 lumenal domain contains six amphipathic helices. Each monomer contributes its α4 and α5 helices, shown as helical wheel plots. (C) Schematic of SigmaR1 domain organization, sites targeted for mutagenesis, and construction of the minimal lumenal domain construct ss-ΔTM-KDEL. (D) Representative images of Cos-7 cells co-transfected with an ER marker (BFP-KDEL) and SigmaR1 point mutants. (E) Quantification of ER sheet area from the experiment in (D) as before. *n* ≥ 26 cells per treatment pooled from 3 experiments. Note, WT control is from Figure 2F-G. Kruskal-Wallis test with Dunn’s multiple comparisons test: **** p < 0.0001, *** p < 0.001, ns p ≥ 0.05. (F) Representative images of Cos-7 cells co-transfected with BFP-KDEL (green) and SigmaR1 amphipathic helix mutants (α4mut is I178A L182A L186A V190A, and α5mut is F196A L199A F200A L203A Y206A L210A L214A Y217A). (G) Quantification of ER sheet area from the experiment in (F). Kruskal-Wallis test with Dunn’s multiple comparisons test: **** p < 0.0001. (H) Representative images of Cos-7 cells co-transfected with BFP-KDEL and SigmaR1 lumenal domain (ss-SigmaR1ΔTM-KDEL) constructs. (I) Quantification of ER sheet area from the experiment in (H). *n* = 30 cells per treatment pooled from 3 experiments. Mann-Whitney test: **** p < 0.0001.

Our goal was to use the details of the molecular structure of the trimeric SigmaR1 complex to inform the mechanism it could use to directly flatten membrane bilayers. Excitingly, the structure of the trimeric SigmaR1 complex^37^ revealed two compelling features which could be ideally suited to membrane flattening: 1) the trimer had a 6.8-nm diameter flat surface that based on the topology would face the lumenal surface of the membrane bilayer and 2) this flat surface had amphipathic helices with hydrophobic residues positioned to further embed the complex within the lumenal leaflet of the bilayer.

First, we sought to determine whether the trimeric state of SigmaR1 is required for its ability to promote ER sheet expansion. We identified two residues that were highly conserved by sequence conservation analysis (Figure S3A) and mediated inter-subunit contacts in the crystal structure that could stabilize the quaternary structure of SigmaR1 (Figure S3B). We mutated each residue individually to alanine (F191A or R119A, Figure 4C-D) and expressed the mutant proteins in Cos-7 cells at levels where wild-type SigmaR1 expanded sheets (Figure 2F-G). We found that the F191A mutant could expand ER sheets to the same degree as wild type, while the R119A mutant expanded ER sheets to a reduced extent (Figure 4D-E). However, when we combined the two mutations together (R119A F191A), SigmaR1 could no longer drive ER sheet expansion (Figure 4D-E). We obtained similar results upon mutating a different conserved binding residue, D188 (Figure S3C-D). We conclude that trimeric assembly of SigmaR1 is an essential feature of its function.

Next, we tested whether the amphipathic nature of the helices facing the membrane bilayer are required for sheet shaping. The flat surface of the SigmaR1 trimer positions a combined six α helices with amphipathic character (from α4 and α5) up against the lumenal surface of the ER membrane bilayer (Figure 4A-B). To test if the membrane-facing hydrophobic residues within these helices are required for SigmaR1 function, we mutated them to alanines. We found that mutating either the α4 or α5 helix alone disrupted the ability of SigmaR1 to drive ER sheet expansion (Figure 4F-G). These results indicate that SigmaR1 requires the amphipathic character of both of its membrane-facing helices to flatten the ER.

Next, we wanted to clarify what role the transmembrane (TM) anchor plays in SigmaR1 function within the lumen of ER sheets. We swapped the short N-terminal tail and TM domain of SigmaR1 with the cleavable signal sequence from the lumenal protein BiP/GRP78 (Figure 4C). With this design, the SigmaR1 lumenal domain is translocated as a soluble protein into the ER lumen. To ensure the retention of this “minimal” SigmaR1 in the ER, we grafted a C-terminal KDEL sequence onto the construct (Figure 4C). We transfected this construct, ss-SigmaR1ΔTM-mNeon-KDEL, into Cos-7 cells along with BFP-KDEL to score its effect on ER localization and morphology. Lumenal SigmaR1 lacking its TM domain localized exclusively to ER sheets, and its expression drove ER sheet expansion (Figure 4H-I). As a control comparison, we confirmed that the lumenal SigmaR1 construct carrying mutations that disrupted its oligomerization (R119A F191A) localized to both sheets and tubules and was unable to expand ER sheets (Figure 4H-I). We conclude that SigmaR1’s lumenal domain is sufficient to partition it into ER sheets and promote sheet expansion.

### SigmaR1 lumenal domain oligomerizes on membranes and inhibits tubule formation

Our data in cells all suggest that lumenal SigmaR1 trimers function by binding and directly flattening the lumenal surface of ER membranes into sheets. To test this model more directly, we purified recombinant SigmaR1 lumenal domain (SigmaR1ΔTM) and performed fluorescence recovery after photobleaching (FRAP) to determine if SigmaR1ΔTM alone was capable of forming a scaffold on synthetic membranes in a simple purified system. Purified SigmaR1ΔTM was dye labelled, incubated with giant unilamellar vesicles (GUVs), and subjected to FRAP. SigmaR1ΔTM bound to the GUVs, indicating that the lumenal domain is sufficient to anchor the complex to membranes (Figure 5A-B). Essentially no recovery was observed for wild-type SigmaR1 over 30 minutes after photobleaching, while full recovery was observed for membrane lipid (Figure 5A-B, E). This indicates that SigmaR1 itself has formed an immobile assembly on the membrane but does not immobilize membrane lipids. The SigmaR1 oligomerization mutant, R119A F191A, in contrast, recovered 54 ± 22% during the same interval (Figure 5C-E). These data are consistent with the in vitro crosslinking assays we performed (Figure 5F-G), which revealed the wild-type SigmaR1ΔTM protein could crosslink into dimers and trimers to a significantly higher degree than the R119A F191A mutant. However, both the crosslinking and FRAP data suggest the mutant is still able to oligomerize at lower levels.

**Figure 5.**
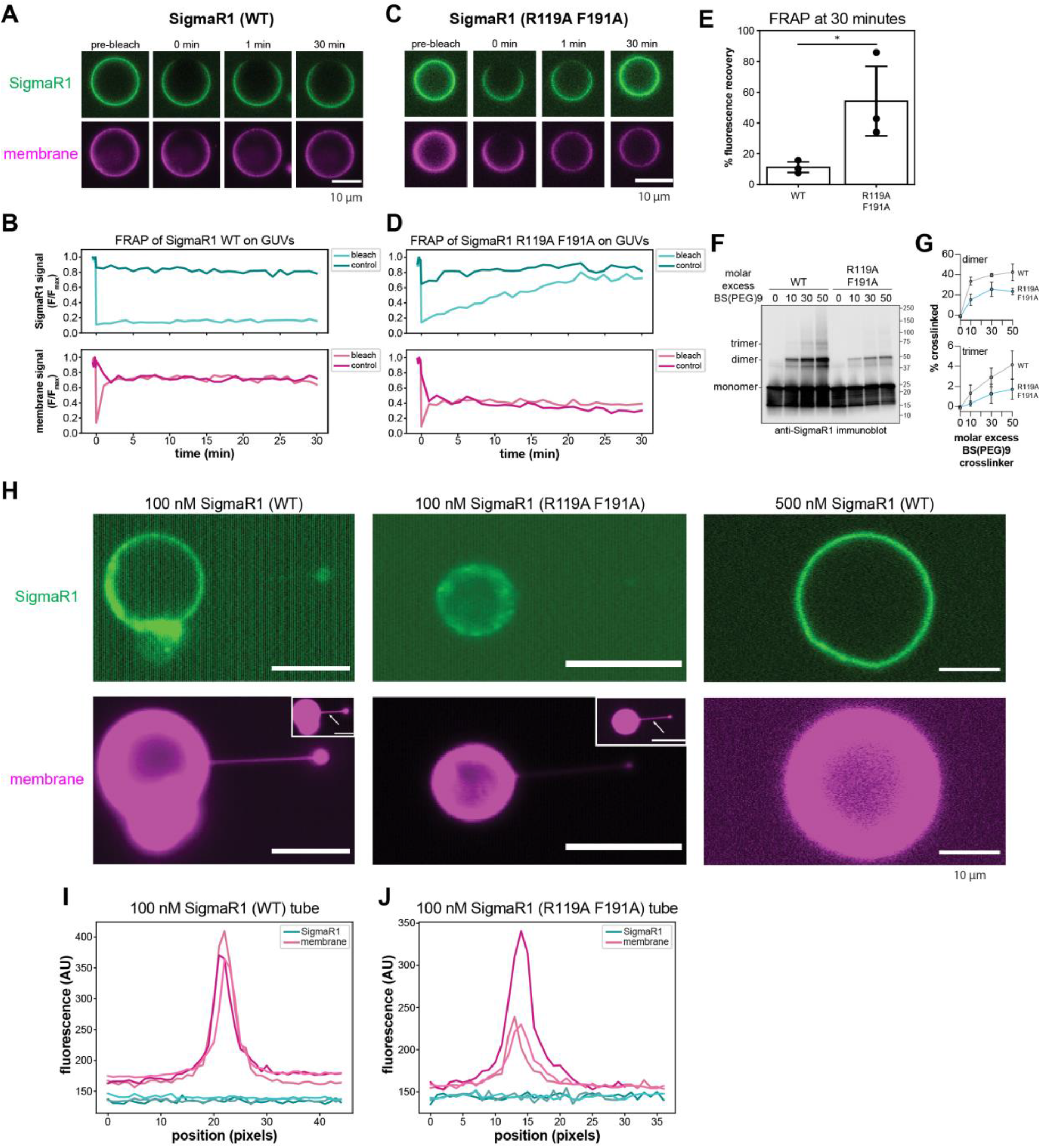
SigmaR1 lumenal domain oligomerizes on membranes in vitro and inhibits tubule formation. (A) Representative images of wild-type SigmaR1ΔTM (green) and GUV membrane (magenta) FRAP. (B) Timecourse of fluorescence signal of a single FRAP event as a fraction of maximum fluorescence value (F/F_max_) for bleached and control regions of wild-type SigmaR1ΔTM sample in (A). Upper plot is protein signal (dark teal: control region, light teal: bleach region). Lower plot is membrane signal (dark violet: control region, light violet: bleach region). (C) Representative images of SigmaR1ΔTM R119A F191A (green) and GUV membrane (magenta) FRAP. (D) Timecourse as in (B) for SigmaR1ΔTM R119A F191A. (E) Bar plot showing percent fluorescence recovery of wild-type (*n* = 3) and R119A F191A mutant (*n* = 3) SigmaR1 protein. One-tailed, equal variance student’s t-test * p < 0.05. (F) Immunoblot of purified recombinant SigmaR1 after treatment with BS(PEG)9 crosslinker in vitro. Monomers and species generated by crosslinks to dimer or trimer are marked. (G) Band intensity quantification of the crosslinked species in (F), mean ± standard deviation, n = 3 independent experiments. (H) Representative confocal fluorescence microscopy images of 100 nM wild-type and mutant SigmaR1ΔTM protein localization on optically trapped membrane tubes pulled from GUVs, showing ATTO488-SigmaR1 (cyan) and ATTO647N-labeled membrane (red). (*n* = 3) Wild-type SigmaR1ΔTM at 500 nM concentration inhibited tube pulling (*n* = 3). (I) Line scans of cross-sections of membrane tubes pulled from GUVs with 100 nM wild-type SigmaR1ΔTM, showing membrane channel signal in magenta and ATTO488 channel signal in teal for *n = 3* replicates. (J) Line scan of membrane tubes as in (I) for 100 nM R119A F191A mutant SigmaR1ΔTM-bound GUVs (*n* = 3). Scale bars: 10 µm.

To test the membrane curvature preference of SigmaR1, we set up experiments using an optical trap to pull membrane nanotubes from GUVs, as in ^41,42^. This experiment uses an optically trapped bead to manipulate the membrane of a GUV to generate a membrane tube of high curvature (∼20 nm radius) contiguous with the GUV, which has relatively flat curvature (∼10 µm radius). This approach has been used to evaluate the membrane curvature sensitivity of membrane binding proteins through comparison of protein density on the membrane surface of the tube to that on the GUV. It would be expected that a protein that preferentially associates with, or stabilizes, flat membranes would be enriched on the GUV surface compared to the membrane tube.

We analyzed confocal fluorescence microscopy images of membrane tubes for relative occupancy of SigmaR1 wild-type and R119A F191A mutant protein on the GUV surface compared to the membrane tube. In the presence of 100 nM wild-type or mutant SigmaR1 (0.2±0.05% coverage of the GUV surface, see Methods), tubes could be pulled from the GUV, but there was no detectable SigmaR1 on the membrane tube for either the wild-type or mutant (Figure 5H-J). At low concentrations, it is expected that unless a protein is enriched on the membrane tube compared to the GUV surface, the protein signal on the tube would be undetectable. When the assay was performed with 500 nM wild-type or mutant SigmaR1 protein, it was not possible to pull membrane tubules from the GUV with the optical trap (Figure 5H, S4). Of note, this protein concentration is within the standard range of the assay and would not be expected to stabilize the GUV via a non-specific effect.^41^ At 500 nM protein, we estimate that SigmaR1 covers 0.9±0.2% of the GUV surface. Although the mutant SigmaR1 was more mobile than wild-type in FRAP (and crosslinking) experiments (Figure 5A-G), the mutant also prevented tubes from being pulled at the higher protein concentration tested. This may be due to the formation of a rigid scaffold *in vitro* by a fraction of the mutant protein, as deduced from FRAP. Collectively, these data show that coverage of just ∼1% of the GUV surface by SigmaR1 is sufficient to destabilize tubule formation.

### SigmaR1 and reticulons are mutually exclusive

The reticulon family of proteins localize to ER tubules.^7^ Rtn4a expands tubular ER when overexpressed, and tubular ER is converted into sheets when Rtn4a is depleted.^7,43,44^ These properties are exactly the opposite of SigmaR1, which localizes exclusively to ER sheets, expands ER sheets when overexpressed, and reduces ER sheets when depleted. Therefore, we asked what happens in the tug of war for peripheral ER morphology when both SigmaR1 and Rtn4a are overexpressed in the same cells. Cos-7 cells were co-transfected with sheet-expanding levels of SigmaR1-mNeon and tubule-expanding levels of Rtn4a-mCherry. We observed a clear separation of SigmaR1-marked ER sheets and Rtn4a-marked ER tubules (Figure 6A). We quantified the degree of SigmaR1 and Rtn4a overlap by the Manders overlap coefficient (Figure 6B), which measures the fraction of SigmaR1 signal overlapping Rtn4a-marked ER regions. To account for random overlap due to ER crowding, we also measured the Manders overlap coefficient in each image with one channel rotated by 90° to randomize pixel distribution. We found a low degree of overlap between SigmaR1 and Rtn4a that nearly matched the overlap calculated using the rotated images (Figure 6B). These data show that SigmaR1 and Rtn4a are unable to localize to the same domains of the ER, and each is capable of maintaining its morphological domain in the presence of the other (see model in Figure 6C). We repeated our Manders overlap analysis using a different reticulon paralog, Rtn3L, that localizes to ER tubules but unlike Rtn4a does not drive tubule expansion.^21^ When scored as before, we observed that SigmaR1 and Rtn3L segregated into non-overlapping domains, and SigmaR1 appeared dominant in its ability to expand ER sheets in the presence of Rtn3L (Figure S5A-B). A significantly different result is observed upon co-expression of Climp63 and Rtn4a.^15,21^ This combination led to a reduction in ER sheet levels and a redistribution of Climp63 into the Rtn4a-labeled ER tubules, which could be scored both visually and quantitatively (Figure 6A-B, Manders overlap coefficient median = 0.86). Together, these results demonstrate that SigmaR1 and reticulon have mutually exclusive activities, and because both are directly shaping the membrane, no membrane shape can accommodate both proteins (Figure 6C). By multiplying the cellular abundances of Rtn4 and SigmaR1 by the dimensions of their molecular footprints on membranes, we estimate that they could cover approximately 0.03% (for Rtn4) to 0.07% (for SigmaR1) of their preferred ER domain (Figure S6). While neither protein is abundant enough to coat membranes, their effects on membrane curvature could be amplified by the crowded environment of cellular membranes and through higher order oligomerization. Interestingly, the placement of SigmaR1 inside the ER is the opposite of reticulons (Figure 6), which assemble on the cytosolic outer leaflet of the ER membrane to generate positive curvature.^7^

The unique structure and topology of SigmaR1 provide a mechanism for rough ER sheet shaping. SigmaR1 trimeric complexes insert six amphipathic helices into the inner leaflet of the ER membrane. These amphipathic helices all lie in a flat plane, likely held rigid by the cupin homology fold that constitutes the majority of the lumenal domain.^37^ This symmetrical, starlike configuration likely cancels out the curvature-inducing effect that inserting a single amphipathic helix would have.^45^ As a result, SigmaR1 trimers partition away from curved membranes and stabilize flat membranes. While many instances of curvature-preferring and curvature-insensitive proteins have been described,^46^ there have been virtually no reports of flat membrane-preferring proteins. Together, our structure-function experiments support a model that the flat amphipathic surface of the SigmaR1 trimer scaffolds flat ER membrane sheets by stabilizing zero curvature within the inner leaflet. SigmaR1 illustrates a new mechanism for how a protein domain can target to and propagate flat membrane sheets from within the lumen.

**Figure 6.**
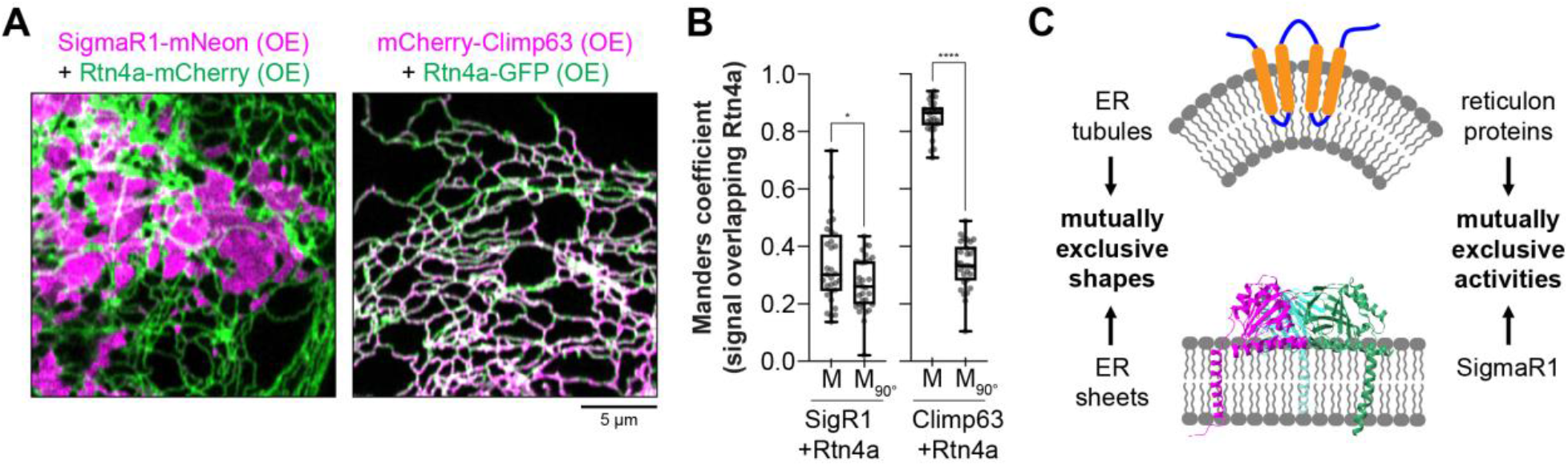
SigmaR1 and Reticulon-4 are mutually exclusive. (A) Representative images of Cos-7 cells co-transfected with Rtn4a-mCh and SigmaR1-mNeon or Rtn4a-GFP and mCh-Climp63. (B) Manders overlap quantification of images taken from the experiment in (A). Analysis was performed with channels in register (“M”) as well as with 1 channel rotated by 90° (“M_90°_”). *n* ≥ 30 cells pooled from 3 experiments. Wilcoxon matched-pairs signed rank test: **** p < 0.0001, * p < 0.05. (C) Model of interplay between SigmaR1 and reticulon proteins: the biochemical activities of SigmaR1 and reticulons are mutually exclusive, as are the underlying membrane morphologies of flat sheets and curved tubules.

## DISCUSSION

We have provided multiple lines of evidence that SigmaR1 functions directly to stabilize ER sheets. SigmaR1 depletion caused a reduction in the level of ER sheets, while its overexpression was sufficient to expand rough ER sheets. Indeed, SigmaR1 can oppose tubule formation and partition into flat membrane sheets in mammalian cells and in vitro. These activities are the opposite of reticulons, which oppose sheet formation and partition into tubules.^7–9^ Thus, ER morphology may be determined by the balance of SigmaR1, reticulons, and membrane lipids. SigmaR1 mutations are linked to juvenile ALS; thus it joins the large cohort of ER-shaping and membrane trafficking proteins implicated in heritable neuronal dysfunction.^31,47^

Interestingly, exogenous ss-SigmaR1ΔTM-mNeon-HDEL protein could even partition to ER sheets and promote ER sheet expansion in a yeast heterologous system (Figure S7A-B). It is notable that although SigmaR1 shows some homology to the fungal sterol isomerase, Erg2, overexpressed yeast Erg2 did not partition into ER sheets or alter ER morphology (Figure S7A-B). Yeast ER morphology may be determined mainly by the ratio of membrane lipids to reticulons.^14^ Although Erg2 and SigmaR1 are structurally similar, the oligomerization state of Erg2 is not known and may be insufficient to stabilize ER sheets. This divergence in function is also consistent with published data that suggest the human sterol isomerase EBP is the functional homolog of Erg2, not SigmaR1.^34–36^ EBP catalyzes the same step in sterol synthesis as Erg2 and unlike SigmaR1 can rescue a yeast *erg2Δ* mutant.^34–36^ ER sheet levels were also unchanged in Cos-7 cells under EBP overexpression, and unlike SigmaR1, EBP localized throughout the ER (Figure S7C-D), indicating that sterol isomerases do not generally alter ER morphology or prefer ER sheets. Finally, we did score whether SigmaR1 overexpression would induce an ER stress response and observe no effect on the three branches of the unfolded protein response (UPR) nor on their normal activation after treatment with the glycosylation inhibitor tunicamycin, Tm (Figure S7E).

In a minimal reconstituted in vitro system, we found that the lumenal domain alone is sufficient to bind membranes and was highly immobile (Figure 5A-E). Higher order SigmaR1 oligomers described in previous studies^37,48–51^ and observed in this study by FRAP and crosslinking likely enhance its membrane-flattening and tubule-destabilizing capacity.^8,45,52,53^ The observed absence of SigmaR1 proteins on tubules pulled in the presence of 100 nM SigmaR1 does not differentiate between a lack of curvature sensitivity vs. a preference for flat membranes. This is a limitation of the study, imposed by the detection limit of the confocal fluorescence experiment. We could not pull tubules out of the GUV in the presence 500 nM SigmaR1 (Figure 5H-J, S4), which corresponds to ∼0.9% coverage of the surface area. This could in principle be due to an increase in the rigidity of the GUV or destabilization of tubule formation. We favor the latter model because it is also consistent with the biological observation that SigmaR1 promotes ER sheet formation relative to tubules even at very low surface density (Figure S6). A high-resolution structure of the SigmaR1 homo-oligomer in the presence of membrane and further biophysical analysis will be required to fully elucidate the physical basis for the ability of SigmaR1 to antagonize tubule formation even at low surface densities.

Lyric and Lrrc59 appear to act through a different, unknown mechanism, not dependent on SigmaR1. We tested whether depletion of SigmaR1 in the backgrounds of Lyric or Lrrc59 overexpression would prevent ER sheet expansion. However, Lyric/Lrrc59 did not need SigmaR1 to promote ER sheet expansion, which indicates that these ER RNA-binding proteins may act through different mechanisms (Figure S8). Lyric and Lrrc59 might play more direct roles in organizing translation on ER sheets, as both proteins associate with mRNAs encoding membrane and secreted proteins^23,24^ and co-precipitate ribosomal proteins and known translocation factors (Figure 2C-D). While neither Lyric nor Lrrc59 contain obvious features that would allow them to stabilize ER sheets, it is possible that experimentally determined structures would provide insight into their sheet-preferring and propagating abilities.

## Supporting information

Script S1

Script S2

## ACKNOWLEDGMENTS

We thank J. Lee, H. Wu, and M. Ricker for reagents and J. Lee for providing unpublished observations about Lyric. We thank C. Ebmeier for performing the mass spectrometry analysis and the Boulder EM Service facility, especially G. Morgan and E. O’Toole for technical advice and assistance. We thank S. Bahmanyar, J. Lee, T. Nguyen, G. Odorizzi, M. Stowell, J. Striepen, and H. Wu for helpful discussions. G.K.V. is an Investigator of the Howard Hughes Medical Institute. We also acknowledge funding from National Institutes of Health Kirschstein NRSA fellowships F32GM137476 (to E.M.S.) and F32AI155226 (to K.P.L.).

## AUTHOR CONTRIBUTIONS

E.M.S., L.E.J., J.H.H., and G.K.V. designed the research plan and interpreted the results. E.M.S., L.E.J., J.B.M., K.P.L., and D.A.P. performed experiments and analyzed the results. E.M.S. wrote the manuscript. G.K.V. and J.H.H. edited and revised the manuscript.

