## Supplemental Information for "A Flat Protein Complex Shapes Rough ER Membrane Sheets"

### Experimental Procedures

Plasmids and siRNAs

This study used the following published plasmids: mNeonGreen-Sec61β,[21] BFP-KDEL,[54] mCherry-Climp63[15] (Addgene #136293), Rtn4a-GFP,[8] Rtn4a-mCherry,[55] Rtn3L-mNeonGreen,[21] and YIPlac204TKC-dsRed-Express2-HDEL (Addgene #21770).

All plasmids generated in this study for transient transfection contained human cDNAs cloned into plasmids mNeonGreen-N1, mNeonGreen-C1, mCherry-N1, or mCherry-C1 (Clontech, Mountain View, CA). Lyric-mNeonGreen was a gift from Jason Lee and contains the human cDNA cloned into the XhoI and BamHI sites of the mNeonGreen-N1 vector. Lyric was subcloned into the XhoI and BamHI sites of mCherry-N1 to generate Lyric-mCherry. Lrrc59 plasmids contain the human cDNA cloned into XhoI and EcoRI sites. SigmaR1 plasmids contain the human cDNA cloned into XhoI and KpnI sites of mNeonGreen-N1 or mCherry-N1. mNeonGreen-EBP was a gift from Molly Ricker and contains the human cDNA cloned into XhoI and BamHI sites. mNeonGreen-KDEL was a gift from Haoxi Wu and contains the signal sequence from BiP/Grp78 (residues 1-18) fused to mNeonGreen with a C-terminal Lys-Asp-Glu-Leu ER retention sequence. pRS423-ss-mNeonGreen-HDEL (2μ plasmid used for yeast overexpression) contains the *TPI1* promoter, the Kar2 signal sequence (amino acids 1-45), mNeonGreen, a C-terminal His-Asp-Glu-Leu sequence, and the *ADH1* terminator from *Candida albicans*, all cloned into the PspOMI and SacII sites. pRS423-ss-SigmaR1ΔTM-mNeonGreen-HDEL was generated by cloning the human SigmaR1 cDNA in between the signal sequence and mNeonGreen. pRS423-Erg2-mNeonGreen contains the *ERG2* promoter region (-1 to -1000 bp) and coding sequence cloned into PspOMI and BamHI sites, followed by in-frame mNeonGreen and the *ADH1* terminator from *Candida albicans* as before. For all constructs, the SigmaR1 lumenal domain (SigmaR1ΔTM) refers to residues 31-223(end). For eukaryotic expression constructs, the lumenal domain was added to the 3' end of the BiP (human) or Kar2 (yeast) signal sequence. For bacterial expression, the SigmaR1 lumenal domain (wild type or R119A F191A mutant) was cloned into pET28-6x His-SUMO (a gift from Greg Odorizzi) using its BamHI and NotI sites. The Ulp1 protease used in the purification was expressed from pET28b-6x His-Ulp1$^{403-621}$ (a gift from Steven Markus).

Lentiviral shuttle vectors were generated by subcloning transient transfection constructs into the pLV backbone (Addgene #36083). Zeocin-selectable Lyric-3x FLAG-mNeonGreen and 3x FLAG-mNeonGreen-Lrrc59 also contained 3x FLAG (added to primers during PCR amplification), an IRES (amplified from Addgene #65278), and BleR drug resistance marker (amplified from Addgene #52961).

Point mutations were generated using the Q5 Site-Directed Mutagenesis Kit (NEB E0554S). The α4 amphipathic helix mutant contained point mutations I178A L182A L186A V190A, and the α5 amphipathic helix mutant contained point mutations F196A L199A F200A L203A Y206A L210A L214A Y217A.

siRNAs were purchased from Dharmacon/Horizon Discovery. Pooled siRNAs targeted Lyric (Dharmacon L-018531-01-0005), Lrrc59 (Dharmacon L-010669-00-0005), or SigmaR1 (Dharmacon L-017475-00-0005). The negative control siRNA was from Thermo Fisher (AM4635). SigmaR1 constructs were rendered siRNA-resistant by replacing the wild-type coding sequence with a version containing silent mutations carried by a dsDNA gBlock (Integrated DNA Technologies).

Cell Culture, Transfection, and Microscopy

Authenticated and mycoplasma tested Cos-7 cells were purchased from ATCC (cat no. CRL-1651, lot #63624240). Authenticated and mycoplasma tested HEK 293T cells were also from ATCC (cat no. CRL-3216, lot #70041165). Authenticated and mycoplasma tested HeLa cells were from ATCC (cat no. CCL-2). Huh-7 cells were a gift from Sara Sawyer. All mammalian cell lines were grown in DMEM (Gibco 12430-062) supplemented with PenStrep (Gibco P4458) and 10% fetal bovine serum (Sigma 12306C), i.e. complete DMEM. Cells were grown at 37°C with 5% $CO_2$.

For microscopy, mammalian cells were seeded into 35-mm imaging dishes with embedded #1.5 cover glass (CellVis D35-20-1.5-N). The next day, cells were transfected with plasmids and lipofectamine 3000 (Invitrogen L3000150) in Opti-MEM I media (Gibco 31985-088) for 5 h at 37°C. Media was replaced with complete DMEM and cells recovered overnight. The next day, cells were washed with FluoroBrite DMEM (Gibco A1896702) supplemented with 10% fetal bovine serum, PenStrep, GlutaMAX (Gibco 35050061), and 25 mM HEPES pH 7.4, i.e. complete FluoroBrite DMEM. Media was replaced after washing with fresh complete FluroBrite DMEM.

Mammalian cells were imaged in a Zeiss LSM880 Airyscan microscope with 405 nm laser for BFP, 488 nm laser for mNeonGreen and GFP, and 561 nm laser for mCherry. The following characteristics were kept constant in all experiments: acquisition with the Airyscan Fast mode and a 63x (1.4 NA) Plan Apo objective, line scanning mode, no line averaging, 0.77 µs pixel dwell time (scan speed 6), super-resolution/SR (2.0x Nyquist) sampling, zoom 1.4 (resulting in 95.97 µm x 95.97 µm or 2716 x 2716 pixel scan area), detector gain 850, and digital gain 1.0 (i.e., no digital gain applied). We used a 488/561/633 nm multiple beam splitter for visible light laser lines and a 405 nm beam splitter for the UV laser line. For two-color imaging of GFP/mNeonGreen and BFP, we used a BP 420-480 + BP 495-550 emission filter. For two-color imaging of mCherry and BFP, we used a BP 420-480 + LP 605 emission filter. For two-color imaging of mCherry and GFP/mNeonGreen, we used a BP 495-550 + LP 570 emission filter. For three-color imaging of mCherry, GFP/mNeonGreen, and BFP, we used the plate setting (no emission filter).

The following concentrations of plasmids were used for transfection of Cos-7 cells (in 2 mL of Opti-MEM I containing transfection reagents): 12.5 ng/mL for low level or rescue expression of SigmaR1, Lyric, and Lrrc59; 75 ng/mL for medium level overexpression of SigmaR1, Lyric, and Lrrc59; 150 ng/mL for high level overexpression of Lyric and Lrrc59 (used only in Fig. 1); 100 ng/mL for overexpression of Sec61β; 25 ng/mL for low level expression of Climp63; 150 ng/mL for overexpression of Climp63; 100 ng/mL for KDEL plasmids; 150 ng/mL for overexpression of Rtn3L; 500 ng/mL for overexpression of Rtn4a; and 75 ng/mL for overexpression of EBP. The following concentrations of plasmids were used for transfection of Huh-7 cells (in 2 mL of Opti-MEM I containing transfection reagents): 150 ng/mL or 250 ng/mL of mNeonGreen-KDEL and 250 ng/mL of SigmaR1-mCherry rescue plasmid.

Yeast cells were *Saccharomyces cerevisiae* W303 and derive from strain UB4785, which contains a wild-type *ADE2* gene to reduce autofluorescence (a gift from Elçin Ünal). All strains used in this study contain YIPlac204TKC-dsRed-Express2-HDEL (Addgene #21770) integrated into the *TRP1* locus by the lithium acetate transformation method. 2µ plasmids were transformed into this background for overexpression experiments. For microscopy, yeast cultures were propagated in SD-His medium with 2% glucose to select for plasmids. Cells were harvested by gentle centrifugation (1900 x g for 2 min) at log phase and analyzed as live mounts on microscope slides. Yeast were imaged with a Yokogawa CSU-W1 SoRa spinning disk confocal, which is built on a Ti2 inverted microscope and contains a Hamamatsu ORCA-FusionBT sCMOS camera (Nikon). In all cases, a 60x (1.42 NA) Plan Apo objective was used with 2.8x SoRa magnification for superresolution. The fluorescence channels were 488 nm excitation (FITC emission filter, BP 525/36) and 561 nm excitation (TRITC emission filter, BP 605/52). Cells were imaged as z stacks with a step size of 0.4 µm and deconvolved with a maximum of 20 iterations of a Richardson-Lucy algorithm in NIS-Elements software (Nikon).

To transfect mammalian cells with siRNAs, cells were seeded into 6-well cell culture plates. The next day, cells were washed with serum- and antibiotic-free DMEM (Gibco 12430-062). Then cells were transfected with 50 pmol of each siRNA and 5 µL of Dharmafect 1 Transformation Reagent (Horizon Discovery T-2001-02) for 6 h, washed with complete DMEM, and allowed to recover overnight. The next day, transfected cells were seeded into 35-mm imaging dishes and grown overnight. The following day, cells were transfected again with 50 pmol of the same siRNA and any applicable plasmids using lipofectamine 3000 and Opti-MEM I but omitting the P3000 reagent provided by the kit. After 5 h, the cells were washed with complete DMEM and grown overnight. Cells were imaged and then trypsinized, washed, and lysed in SDS sample buffer (2% SDS, 80 mM Tris pH 6.5, 10% glycerol, 2.5% 2-mercaptoethanol) to obtain a matched protein extract for immunoblot analysis.

For the ER stress experiment, tunicamycin (Sigma T-7765) was added to cells 1 day after transfection for 4 h at a final concentration of 2 µg/mL (from a 2 mg/mL stock).

For production of lentiviral particles, HEK 293T cells were seeded into 6-cm cell culture dishes. The next day, cells were transfected with lipofectamine 3000 and 4.5 µg of pLV shuttle vector, 4 µg of pMDLg-RRE (Addgene #12251), 2 µg of pRSV-REV (AddGene #12253), and 2 µg of pCMV-VSV-G (Addgene #8454) in Opti-MEM I for 3 h. Cells were washed with complete DMEM and viral particles were collected from the culture supernatant after 48 h and 72 h, pooled, and concentrated by centrifugation at 20,000 x g in 1-mL aliquots. For infection, Cos-7 or HeLa cells were seeded into 6-well cell culture dishes. The next day, the media was replaced with complete DMEM containing resuspended lentiviral particles and 10 µg/mL polybrene. Cells were infected overnight, then media was replaced the next day. Cos-7 SigmaR1-mNeonGreen stable overexpression cells were purified by FACS. HeLa cell lines used for immunoprecipitation experiments were selected by 100 μg/mL Zeocin (Thermo R25001) beginning 72 h after initial infection and ending ~14 days after infection when populations were verified pure by microscopy.

Protein Gels, Immunoblotting, and Antibodies

Cell pellets were directly lysed in 1x SDS sample buffer (2% SDS, 80 mM Tris pH 6.5, 10% glycerol, 2.5% 2-mercaptoethanol) and denatured by heating at 50°C for 5 min. Other samples were supplemented with 5x SDS sample buffer to the same final 1x concentration prior to denaturation. Denatured lysates were loaded with a Precision Plus Protein™ All Blue Prestained Protein Standards ladder (Biorad 1610373) on Criterion TGX 4-20% polyacrylamide gradient gels (BioRad 5671094) and run at 200 V for 35 min.

For total protein staining, SDS-PAGE gels were stained with the Colloidal Blue Staining Kit (Thermo LC6025).

For immunoblot, proteins were transferred to Immobilon-P PVDF membrane (Millipore IPVH00010) at 50 V for 1 h. Membranes were blocked with 5% nonfat milk in TBST (block) for 30 min, then incubated with primary antibodies in block overnight at 4°C. The next day, blots were washed 4 times (5 min each) with TBST and incubated with secondary antibodies in block for 1 h at room temperature. Membranes were exposed to SuperSignal West Pico PLUS Chemiluminescent Substrate (Thermo 34580) or SuperSignal West Femto Maximum Sensitivity Substrate (Thermo 34096) reagents and imaged with a ChemiDoc XRS+ System (BioRad).

The following primary antibodies were used: rabbit anti-Lyric 1:750 (Sigma HPA010932), rabbit anti-Lrrc59 1:5000 (Bethyl/Thermo A305-076A), rabbit anti-GAPDH 1:10,000 (Sigma G9545), rabbit anti-FLAG 1:2000 (Sigma F7425), rabbit anti-RPS24 1:2000 (Abcam ab196652), rabbit anti-SIGMAR1 1:1000 (Cell Signaling Technology 61994S), rabbit anti-BiP 1:1000 (Cell Signaling Technology 3177S), rabbit anti-P-eIF2α (phospho-Ser51) 1:1000 (Cell Signaling Technology 3398P), and mouse anti-Atf6 1:1000 (Abcam ab122897). The secondary antibody was goat anti-rabbit HRP conjugate 1:6000 (Sigma A6154).

Immunoprecipitation and Mass Spectrometry

Cells grown in 10-cm cell culture dishes were trypsinized, washed, and collected by centrifugation at 200 x g for 2 min. We used a published protocol as the basis for our immunoprecipitations.[56] Cells were lysed in solubilization buffer (50 mM HEPES pH 7.4, 200 mM NaCl, 2 mM $Mg(OAc)_2$, 1% digitonin [high purity; Calbiochem 300410], and 1:500 Protease Inhibitor Cocktail Set III, EDTA-Free [Calbiochem 539134]) on ice for 30 min with gentle agitation every 10 min. Meanwhile, beads were equilibrated in wash buffer (50 mM HEPES pH 7.4, 200 mM NaCl, 2 mM $Mg(OAc)_2$, 0.1% digitonin). 20 μL of packed anti-FLAG M2 agarose beads (Sigma A2220) were used for each 100 μg of wet cell weight. Solubilized lysates were cleared by centrifugation at 20,000 x g for 10 min at 4°C, then beads were added to the soluble fraction. Tubes were agitated with end-over-end rotation at 4°C for 90 min. Then beads were washed 5 times with 950 μL of wash buffer (incubating for 5 min with end-over-end rotation for the first two washes). Finally, the sample was eluted with a competitor 3x FLAG peptide (Sigma F4799) in elution buffer (50 mM HEPES pH 7.4, 150 mM NaCl, 2 mM $Mg(OAc)_2$, 0.25% digitonin, 250 μg/mL 3x FLAG peptide) for 30 min at room temperature with agitation. Matched samples were analyzed by immunoblot, colloidal Coomassie staining, and LC-MS/MS.

Mass spectrometry on tryptic peptides was performed by the University of Colorado Boulder Mass Spectrometry Facility on an Orbitrap Q-Exactive HF-X LC-MS/MS system (Thermo Scientific). Protein abundances in the starting mixtures were estimated using the label-free iBAQ quantification algorithm in MaxQuant using the human proteome (Uniprot Proteome ID UP000005640). Known contaminants were filtered out from the hits, which were further filtered by a 10-fold enrichment cutoff relative to the untagged control prior to plotting the data.

Fluorescence Image Analysis

Zeiss LSM880 Airyscan images were prepared by the Airyscan Processing function in Zen Black software with default, standard settings. Linear adjustments to brightness and contrast were made in FIJI[57] for image presentation. All quantifications were performed on regions of interest (ROIs) containing resolvable (2D) ER. To reduce bias, ROIs were selected from the north (top) position of the cell when in focus using a standardized procedure.

ER sheet area was quantified using MATLAB script S1. For this analysis, it is important to have high contrast and high resolution images. The MATLAB program generates a binary ER mask using Otsu's threshold method. Then the mask is eroded by the diameter of ER tubules determined empirically from the pixel size of the images. Pixels that persisted after erosion are classified as ER sheet pixels. The % ER sheet area is quantified by dividing the number of ER sheet pixels by the total number of ER pixels. We found that the accuracy of thresholding was improved by including extracellular regions. For analysis of yeast ER, individual cells were manually traced and cropped from original images containing many cells. For accurate segmentation, we found that it was necessary to set the background pixel intensity outside the traced cell but within the square image area to match the background within the cell using the Math > Set function in FIJI.

We quantified Manders overlap using MATLAB script S2. The script generates binary masks of both channels using Otsu's method. The Manders overlaps reported are calculated as the fraction of above-threshold Climp63 or SigmaR1 signal that overlaps the reticulon binary mask. For each image, the analysis is repeated after rotation of one channel by 90°. To ensure accuracy, ROIs did not contain extracellular pixels.

Protein Conservation and Structure Analysis

SigmaR1 amino acid conservation was scored using the ConSurf server[58] based on multiple sequence alignment of 66 homologs. SigmaR1 structures were analyzed and rendered into figures with UCSF ChimeraX.[59] Helical wheel plots were generated using HeliQuest.[60]

Electron Tomography

Cos-7 empty vector control or SigmaR1-mNeonGreen stable overexpression cells were grown on UV-sterilized sapphire discs (Technotrade International 405-300) coated with fibronectin (Sigma F0895). Discs were dipped into cryoprotectant (complete DMEM supplemented with 2% sucrose and 150 mM mannitol) and high pressure frozen in a Wohlwend Compact 02 freezer (Technotrade International). Samples were placed in vials containing acetone, 2% osmium, 0.2% uranyl acetate, and 1% water. After 6 h at -90°C, the vials were slowly warmed to room temperature and infiltrated with epon resin. Discs were then placed on glass slides, covered with a thin layer of resin, and polymerized. Resin-embedded cells were excised and glued to blank resin stubs for sectioning. Thick serial sections (250 nm) were collected and post stained with 2% aqueous uranyl acetate and Reynolds lead citrate. Gold fiducials (15 nm, Ted Pella 15704-20) were applied after staining. Single axis tilt series were collected on a Tecnai F20 electron microscope with a Gatan K3 (4k) direct detection camera or on a Tecnai F30 electron microscope equipped with a Gatan OneView IS (4k) CCD camera.

Tomograms were reconstructed from tilt series using IMOD software.[61] To quantify rough ER profile lengths, two tomographic volumes were extracted from each cell (2060 x 2060 x 343 nm) for a total of 8 volumes per condition. Rough ER membrane profiles were traced on a total of nine Z planes of each volume. Contour lengths were summed and graphed. The profile lengths report all ER membrane lengths in a total of ~76 $\mu m^2$ of sampled 2D regions per cell.

Protein Purification for GUV Experiment

Log-phase *E. coli* C41(DE3) cultures (1 L of LB per purification) carrying 6x His-SUMO-SigmaR1$^{\Delta 1\text{-}30}$ plasmids were induced with 500 μM IPTG at 18°C shaking overnight. The next day, cells were harvested by centrifugation and lysed in His tag lysis buffer (20 mM HEPES-KOH pH 7.4, 300 mM NaCl, 5 mM 2-mercaptoethanol, 1:4000 dilution Benzonase nuclease [Millipore Sigma E1014], 1x cOmplete mini EDTA free protease inhibitor cocktail [Roche 42484600], 5 mM imidazole, and 4% CHAPS [Anatrace C316]). Cells were lysed by incubation at 4˚C with rocking for 30 minutes followed by tip sonication (2 s on, 1 s off, for 10 minutes) in Misonix Sonicator 3000. Insoluble material was cleared by centrifugation at 40,000 x g in a JA-25.5 rotor for 30 min at 4°C. The supernatant was then applied to a 1ml packed, equilibrated TALON metal affinity resin column (Takara 635502).

Beads were washed with 40 ml of His Tag wash buffer (20 mM HEPES-KOH pH 7.4, 300 mM NaCl, 5 mM 2-mercaptoethanol, 5 mM imidazole, 0.4% CHAPS). We eluted the protein by digestion with the SUMO hydrolase, 6x His-Ulp1, which was purified in house. First beads were washed with digestion buffer (20 mM HEPES-KOH pH 7.4, 150 mM NaCl, 0.4% CHAPS). Then beads were incubated with 1 mL of digestion buffer containing 30 µg of 6x His-Ulp1 overnight at 4°C with rocking.

The next day, flowthrough was collected containing processed SigmaR1 lumenal domain. Samples were diluted 1:10 in imaging buffer (20 mM HEPES-KOH pH 7.4, 150 mM NaCl, 5 mM 2-mercaptoethanol) to which was added Bio-Beads (1g per 10 mL of diluted protein eluate) (Bio-Rad 1523920). SigmaR1ΔTM was incubated at 4˚C with rocking for 2 hours. SigmaR1ΔTM was decanted from the Bio-Bead slurry and concentrated to 0.5-1 mg/mL in a 10 kD molecular weight cutoff spin concentrator (Millipore Sigma UFC901024). SigmaR1ΔTM concentration was evaluated by absorbance at 280 nm. Fluorescence labeling was performed by incubating protein with ATTO488 NHS Ester dye at a molar ratio of 1:2 protein to dye, at 4˚C overnight with rocking. Excess dye was removed by passing the labeling reaction over a PD MiniTrap desalting column (Cytiva 28918007). Labeled SigmaR1ΔTM was concentrated in a 10k MWCO spin concentrator to a final concentration of 1 mg/mL, and labeling efficiency was assessed by absorbance at 280 and 488 nm. Protein was stored at 4˚C.

GUV FRAP Experiment

Giant unilamellar vesicles (GUVs) were formed by hydrating a lipid film of 50 nanomoles of lipid of molar composition (70% DOPC, 20% DOPE, 5% DOPS, 5% cholesterol, 0.2% Atto-647N DOPE, and 0.01% 1,2-distearoyl-sn-glycero-3-phosphoethanolamine-N-[biotinyl(polyethylene glycol)-2000] (DSPE-PEG(2000) Biotin)) in 100 µl of 320 mosM sucrose on a polyvinyl alcohol (PVA) support for 30 minutes at room temperature. 500 nM SigmaR1ΔTM was mixed with 5 µl of GUVs in 150 µl of imaging buffer. Fluorescence imaging was performed on a Nikon Ti2 microscope with a Nikon A1 confocal unit. Fluorescence recovery after photobleaching (FRAP) was performed using Nikon Elements software. GUVs were imaged for five frames over 30 seconds before bleaching a region at the top of the GUV. After photobleaching, fluorescence intensity was monitored in the bleached region of the GUV as well as in an unbleached, control region of the same GUV.

GUV FRAP Image Analysis

Median-filtered image background was subtracted from images using FIJI. Mean values of bleached and control regions for each timepoint were calculated using a home-written Python script (https://github.com/livjensen7/GUVscripts/blob/main/GUV_FRAP_Analysis-.ipynb). The percent fluorescence recovery for each bleach region was calculated as the difference of the mean value of the last 4 frames of the bleached region and the mean value of the bleached region immediately after bleaching, all divided by the difference of the mean value of the last 4 frames of the control region and the mean value of the control region immediately after bleaching, i.e.:

$$\frac{F_{final,bleach}-F_{initial,bleach}}{F_{final,control}-F_{initial,control}}.$$

Statistical significance was evaluated by a one-tailed, equal variance student's t-test.

Crosslinking Experiment

SigmaR1ΔTM was purified as described above, with minor modifications. The lysis buffer additionally contained 1 mg/mL hen egg white lysozyme (Sigma L4919), and cells were lysed by incubation on ice for 30 min (with agitation every 10 min) followed by tip sonication (20 s on, ≥ 60 s resting on ice, 20 s on, in a Sonicator Cell Disruptor model W185F [Heat Systems-Ultrasonics, Inc., Plainview, NJ]). The spin-cleared lysate was additionally cleared by filtration through a 0.2 µm filter. For bead capture, lysates were incubated with 300 µL of packed, equilibrated TALON metal affinity resin (Takara 635502) for at least 1 h at 4°C with end-over-end rotation. The wash buffer contained 0.1% CHAPS, and the elution buffer contained 4% CHAPS. No adsorptive detergent removal was performed, and proteins were dialyzed against PBS without dye labeling.

Crosslinking reactions were set up in PBS (Sigma D8537) and contained 3 µM recombinant SigmaR1 and BS(PEG)9 crosslinker (Thermo Scientific 21582) at molar ratios that ranged from 0 (control) to 50-fold excess relative to SigmaR1. Crosslinking samples were incubated on ice for 2 h. Then the reactions were quenched with 20 mM Tris-HCl (pH 7.5) for 15 min and denatured for immunoblotting as described above. Band intensities were quantified in FIJI with local background subtraction.

Membrane Surface Density Estimation

The density of SigmaR1 protein on the GUV surface was estimated for the GUV FRAP experiments as in [41]. The footprint area of a SigmaR1 trimer was estimated to be ~20 $nm^2$ from the crystal structure of SigmaR1 (PDB ID: 5HK1), assuming the footprint was an equilateral triangle of side length ~6.8 nm. To approximate surface density of ATTO488 labeled protein on the GUV surface, a standard curve was generated from quantification of fluorescence intensity of GUVs containing variable amounts of ATTO488-DOPE, ranging from 0.01-0.1 mol %. Surface density of ATTO488-DOPE was calculated using an approximate surface area of 0.7 $nm^2$ per lipid.

Membrane Tube-Pulling Experiments

Tube pulling experiments were performed as in [41] on a Nikon A1 confocal microscope fitted with an optical trap. Briefly, GUVs containing a minor fraction of biotinylated lipids were incubated with 100 nM or 500 nM ATTO488-labeled SigmaR1 protein and added to imaging chambers that had been passivated with 1 mg/ml BSA in imaging buffer. Membrane tubes were formed by bringing a streptavidin coated bead into contact with GUVs which had settled on the surface of the imaging chamber with an optical trap, and then retracting the bead. Protein association with the membrane tube was assessed by quantification of confocal microscopy images.

Graphs and Statistics

All graphs were generated in Prism 9 (GraphPad). Statistical analyses were also performed in Prism 9 (GraphPad). Statistical test designs and results are indicated in the figures and figure legends.

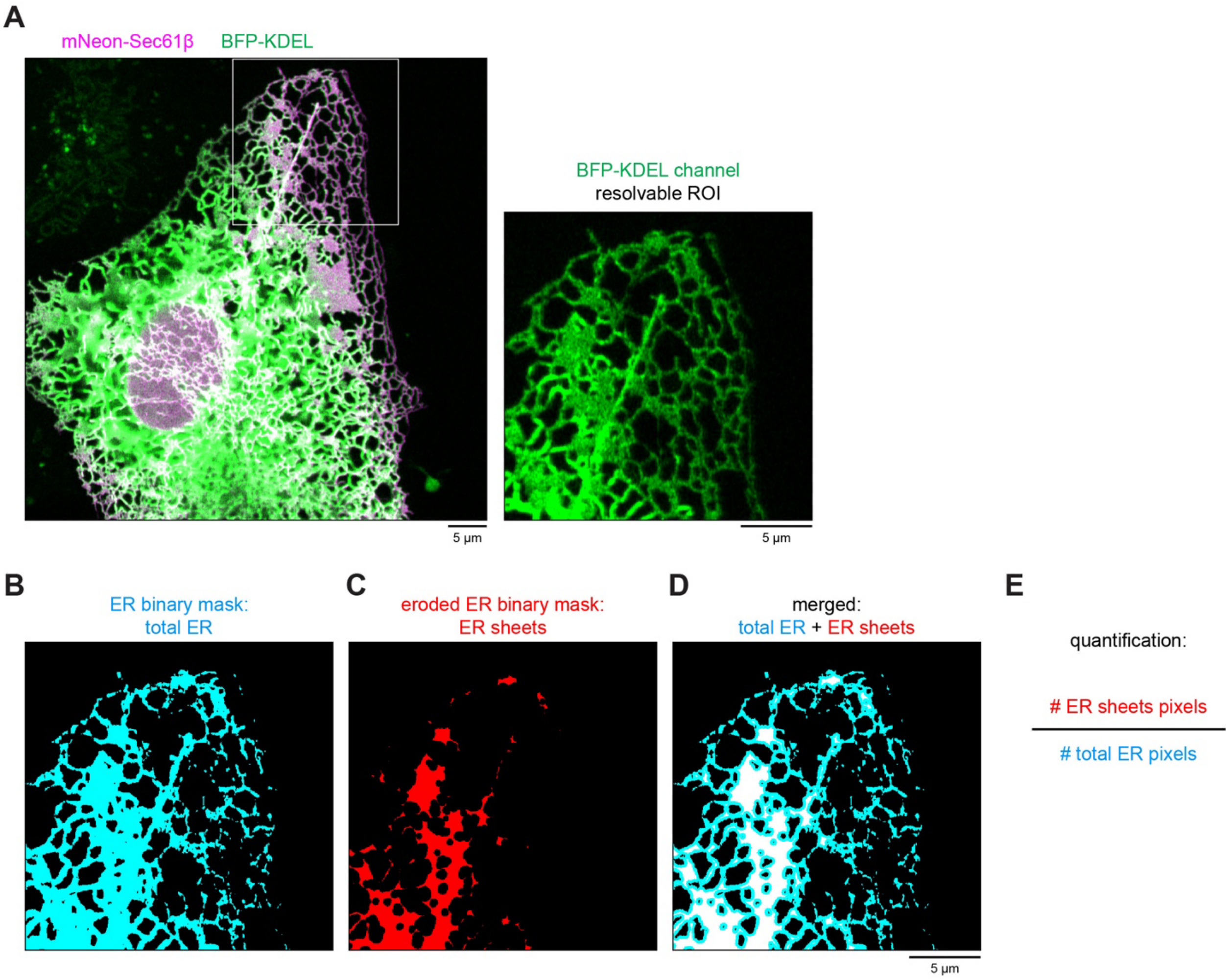


**Figure S1. ER sheet area quantification method.**

(A) Image of a control cell from the experiment in Fig. 1A with example resolvable region of interest from the general marker BFP-KDEL channel.

(B) Binary ER mask generated by Otsu threshold.

(C) Eroded mask showing ER sheets (red).

(D) Merged image of the eroded ER sheet mask over the total ER mask (sheets after merging are white).

(E) Quantification method to calculate % ER sheet area.

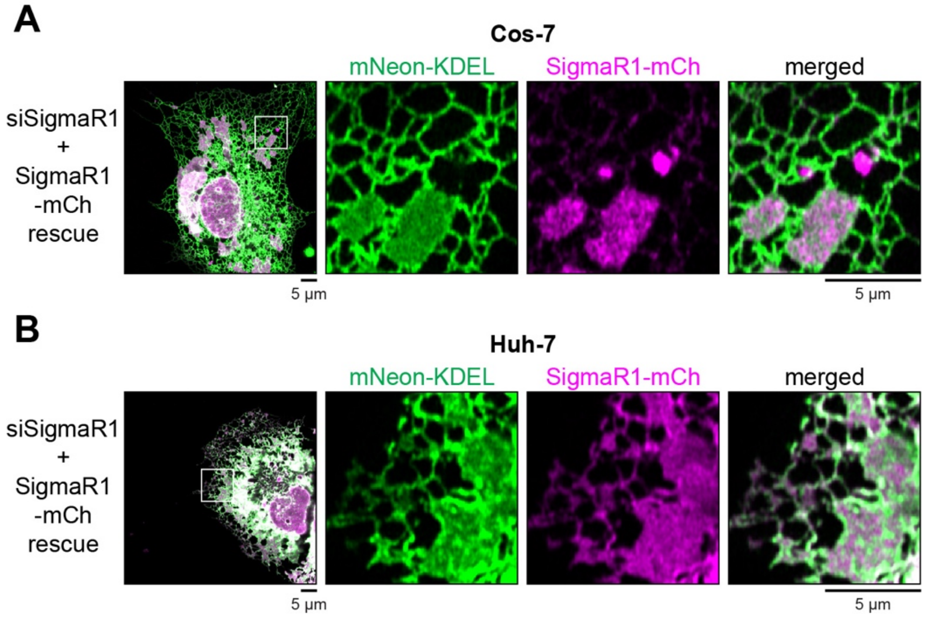


**Figure S2. SigmaR1 rescues ER sheet levels.**
(A) Cos-7 cell depicted in Fig. 2J with mNeon-KDEL and SigmaR1-mCh rescue channels shown.
(B) Huh-7 cell depicted in Fig. 2K with mNeon-KDEL and SigmaR1-mCh rescue channels shown.

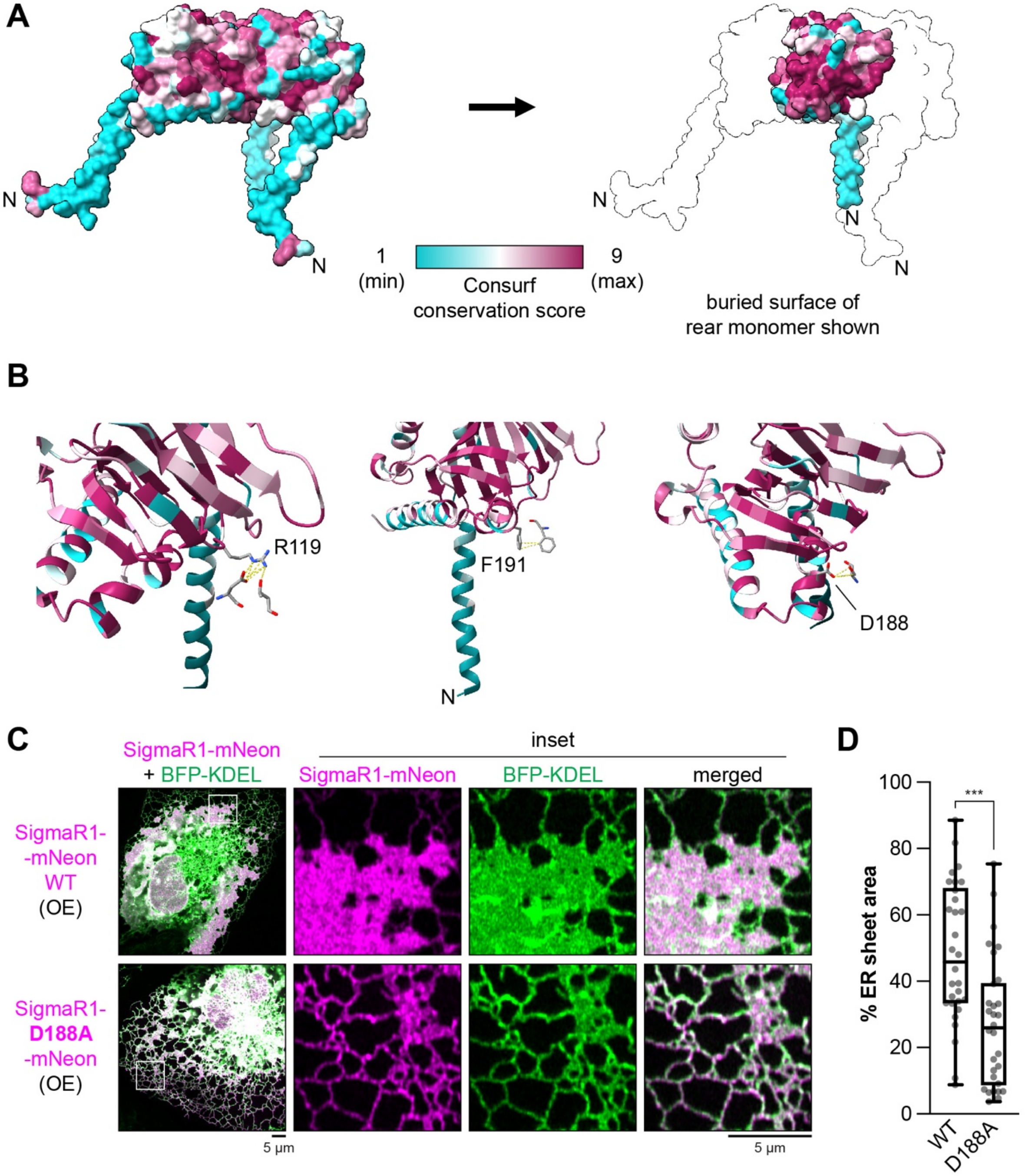


**Figure S3. SigmaR1 trimer assembly is required for ER sheet propagation.**
(A) Surface of SigmaR1 trimer crystal structure (PDB: 5hk1) color-coded by Consurf amino acid conservation score. The buried surface of one monomer is also shown after stripping away the other two monomers.
(B) SigmaR1 crystal structure cartoon of one monomer with conserved binding residues highlighted (R119, F191, or D188) showing cross-subunit contacts.
(C) Representative images of Cos-7 cells transfected with wild-type SigmaR1-mNeon or D188A mutant.
(D) Quantification of ER sheet area from the experiment in (C). *n* = 30 cells per treatment pooled from 3 experiments. Mann-Whitney test: *** $p < 0.001$.

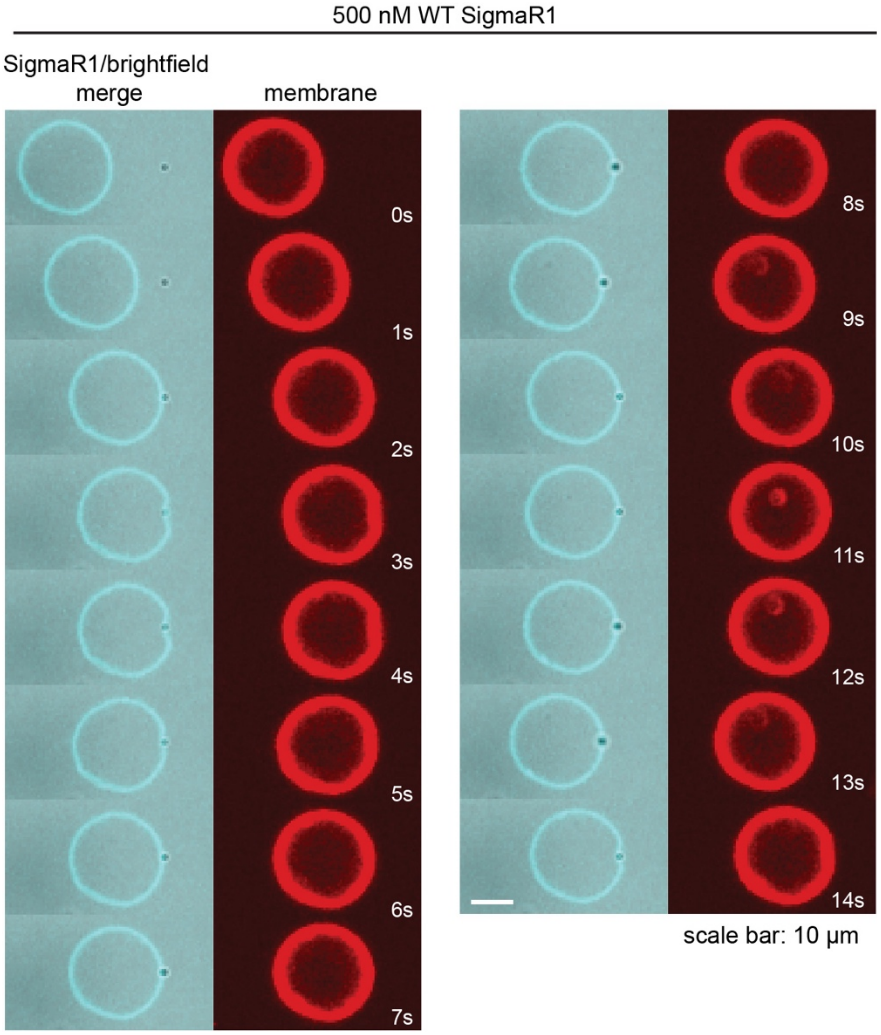


**Figure S4. SigmaR1 inhibits tube pulling.**
Representative confocal fluorescence microscopy time course of a GUV (ATTO647N-labeled membrane, red) with 500 nM SigmaR1 (ATTO488-labeled, cyan, and merged with brightfield) that resists tube pulling by an optically trapped bead. Representative of 3 experiments. Scale bars: 10 µm.

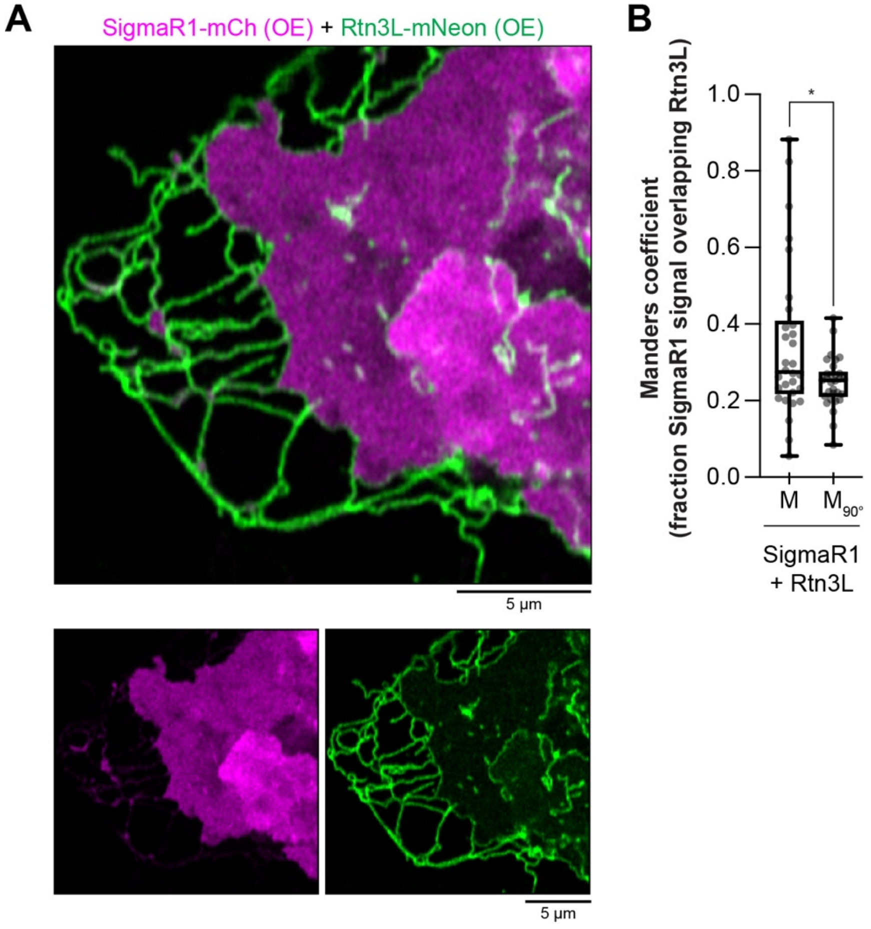


**Figure S5. SigmaR1 ER sheets exclude Rtn3L.**
(A) Representative image of a Cos-7 cell overexpressing SigmaR1-mCh at sheet-inducing levels while simultaneously overexpressing Rtn3L-mNeon.
(B) Manders overlap quantification of the experiment in (A). Analysis was performed with channels in register ("M") as well as with 1 channel rotated by 90° ("$M_{90°}$"). *n* = 30 cells pooled from 3 experiments. Wilcoxon matched-pairs signed rank test: * $p < 0.05$.

| protein | number of molecules per cell | membrane surface covered by each molecule | total membrane surface covered per cell | ER domain area per cell | ER domain coverage |
|---|---|---|---|---|---|
| SigmaR1 | $1\times10^5$ * | 20 $nm^2$ † | 2.0 $\mu m^2$ | 2800 $\mu m^2$ (sheets) | 0.07% |
| Rtn4 | $3.4\times10^5$ | 5.01 $nm^2$ ‡ | 1.7 $\mu m^2$ | 5070 $\mu m^2$ (tubules) | 0.03% |
| Sec61α | $4.7\times10^5$ | 9.6 $nm^2$ § | 4.5 $\mu m^2$ | 7870 $\mu m^2$ (sheets + tubules) | 0.06% |

**Figure S6. Mammalian ER coverage by membrane-shaping proteins.**
Calculation of the ER surface that could be covered by SigmaR1 and Rtn4, with Sec61α as a comparison. ER domain areas from [4] and protein copies per cell from [62]. * Database value divided by 3 to correspond to SigmaR1 trimers. † Area of a triangle bounding the complex from PDB:5HK1 ‡ Area of a rectangle bounding the reticulon homology domain from S. cerevisiae Yop1 (AlphaFold/UniProt:Q12402) § Area of a circle bounding the transmembrane domains from PDB:6W6L

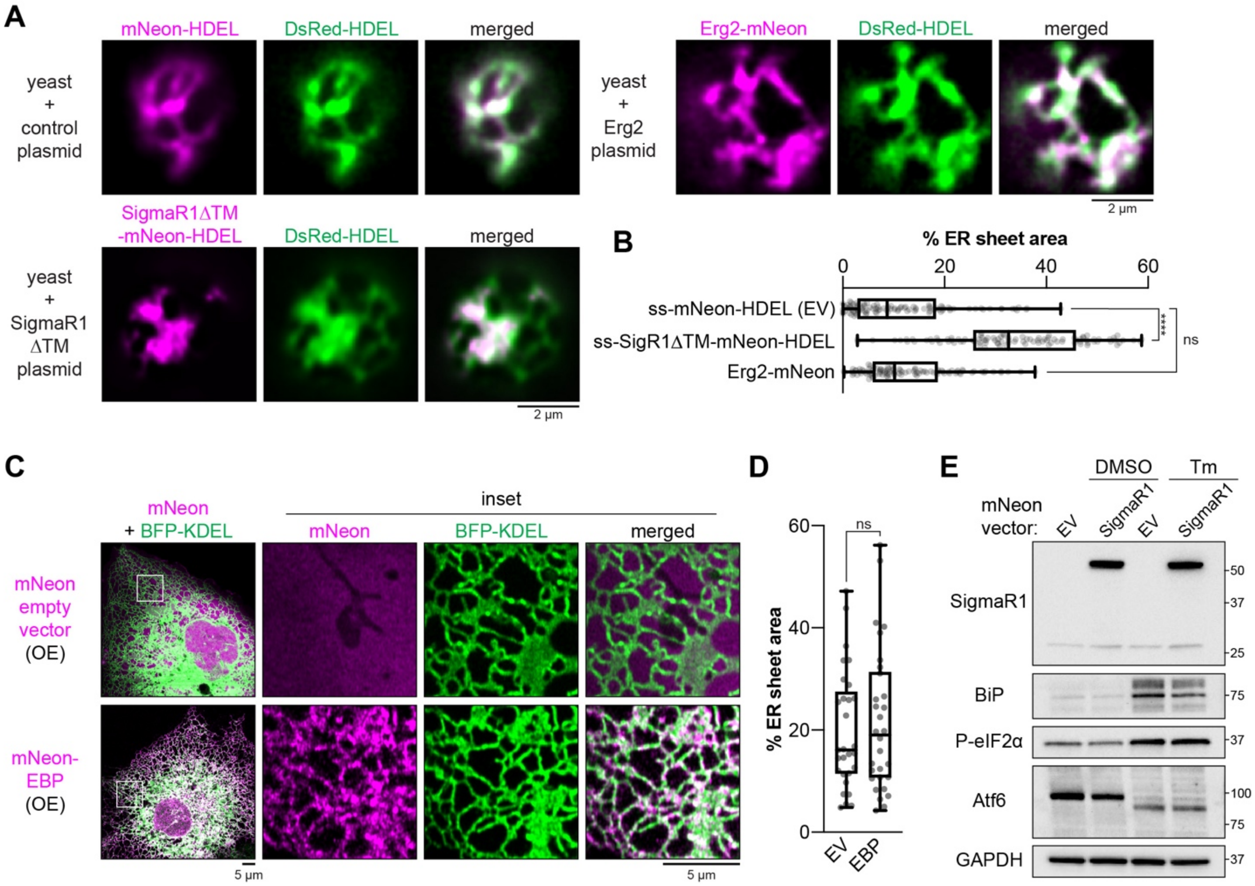


**Figure S7. SigmaR1 function has diverged from sterol isomerases and does not induce ER stress.**
(A) Representative images of ER morphology in budding yeast expressing an ER lumenal marker (DsRed-HDEL, green) and the SigmaR1 lumenal domain (SigmaR1ΔTM-mNeon-HDEL), Erg2 (Erg2-mNeon), or a control (mNeonGreen-HDEL) from a 2µ plasmid.
(B) Quantification of ER sheet area from the experiment in (A). $n = 100$ cells per treatment pooled from 3 experiments. Kruskal-Wallis test with Dunn's multiple comparisons test of each condition compared to the control: **** $p < 0.0001$, ns $p \geq 0.05$.
(C) Representative images of Cos-7 cells overexpressing mNeon empty vector or mNeon-EBP as well as ER marker BFP-KDEL.
(D) Quantification of ER sheet area from the experiment in (C). $n \geq 29$ cells per treatment pooled from 3 experiments. Mann-Whitney test: ns $p \geq 0.05$.
(E) Immunoblot analysis of Cos-7 cells transfected with mNeon empty vector or SigmaR1-mNeon and treated with DMSO control or 2 µg/mL tunicamycin (Tm) for 4 h. Transfection conditions match the higher level SigmaR1 overexpression in Fig. 2F. Membranes were probed with anti-SigmaR1, anti-BiP (reporter for IRE1 pathway), anti-P-eIF2α (reporter for PERK pathway), and anti-Atf6 (cleavage reports activation of Atf6 pathway) antibodies. Representative of 3 biological replicates.

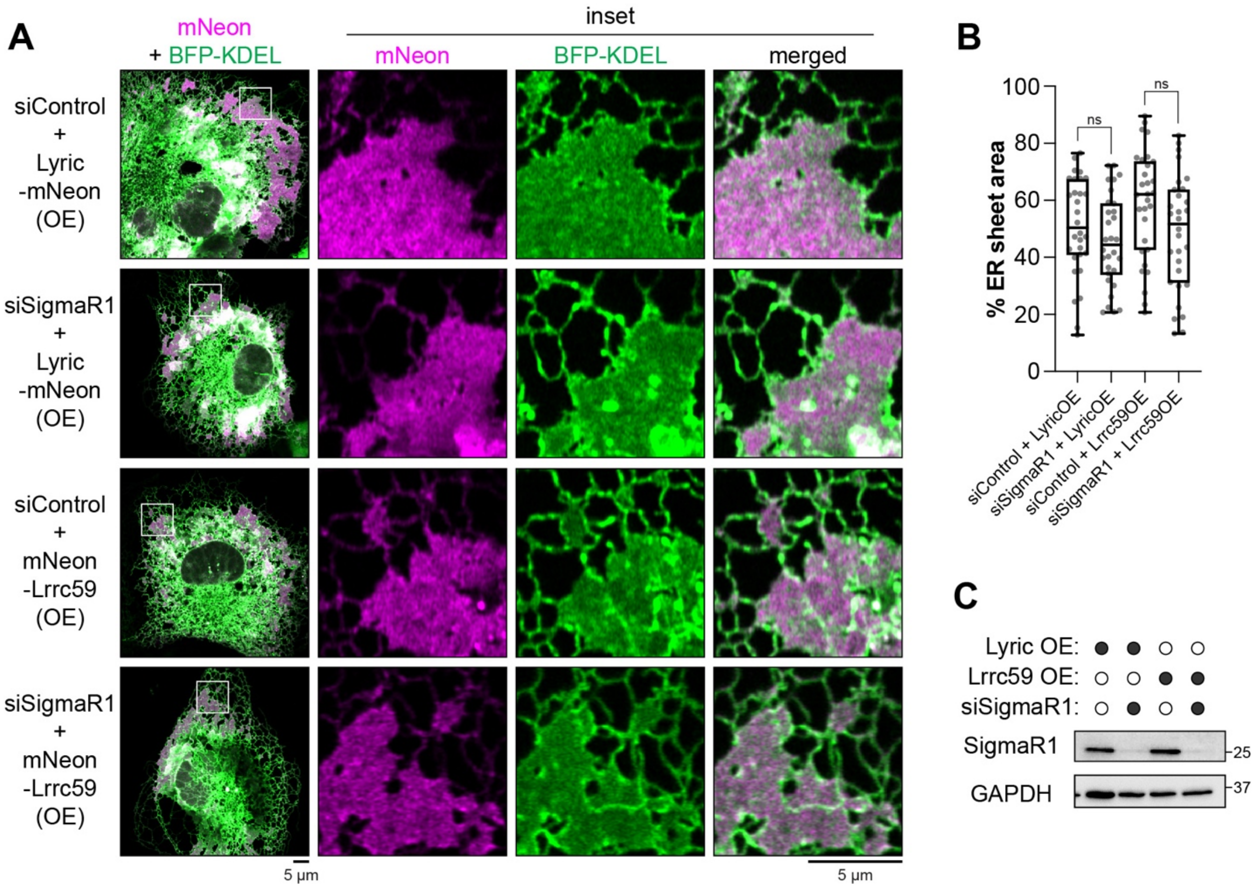


**Figure S8. Lyric and Lrrc59 do not require SigmaR1 to expand ER sheets.**

(A) Representative images of ER morphology in Cos-7 cells co-transfected with control or SigmaR1 siRNAs, Lyric-mNeon or mNeon-Lrrc59 overexpression plasmids, and ER marker BFP-KDEL.
(B) Quantification of ER sheet area from the experiment in (A). *n* = 30 cells per treatment pooled from 3 experiments. Kruskal-Wallis test with Dunn's multiple comparisons test: ns $p \geq 0.05$.
(C) Immunoblot analysis (anti-SigmaR1) of the experiment in (A).

**Movie S1. Electron tomography of empty vector control.** Movie of reconstructed volume of the Cos-7 empty vector control cell in Fig. 3A. Movie plays at 20 frames per second forward and then reverse.

**Movie S2. Electron tomography of SigmaR1-mNeon overexpressing cell.** Movie of reconstructed volume of the Cos-7 SigmaR1-mNeon overexpressing cell in Fig. 3A. Movie plays at 20 frames per second forward and then reverse.
