## Supplementary material for "A Flat Protein Complex Shapes Rough ER Membrane Sheets": Script S1

% calculate_ER_sheet_area.m

% automatically Otsu thresholds images, identifies tubules and sheet
% edges by binary erosion, allowing segmentation of two categories:
% 
% (1) tubules and sheet edges (high curvature)
% (2) sheet faces (low curvature)
%
% returns:
% image file name
% total image area
% total binary ER area
% area of binary ER that is tubules and sheet edges
% area of binary ER that is sheet faces

% requires a vertical cell array of strings, which specifies the file names,
% to be called inputfilenames

% Notes to the User:
% * MATLAB must have the Image Processing Toolbox installed.
% * Images must have high contrast for the segmentation to be accurate.
% * Images must be cropped to ROIs containing only resolvable (2D) ER; 
%   otherwise, crowded ER will be misclassified as ER sheets.
% * Comment/uncomment the final "optional" section depending on if you 
%   want MATLAB to show image processing intermediates. It is useful to 
%   spot check a few images prior to analyzing a batch to verify quality.
% * The variable scalefactor needs to be determined carefully based on 
%   the resolution (pixel size) of the images so that the program erodes 
%   only by the size of ER tubules.

inputfilenames = {'myfile1.tif'
'myfile2.tif'
'myfile3.tif'};

numfiles = length(inputfilenames);

% Generates an output cell array
outputdata = cell(numfiles + 1, 5); % +1 to allow space for header line
outputdata{1, 1} = 'Image';
outputdata{1, 2} = 'Image Area (pixels)';
outputdata{1, 3} = 'ER Area (pixels)';
outputdata{1, 4} = 'Tubule and Sheet Edge Area (pixels)';
outputdata{1, 5} = 'Sheet Face Area (pixels)';

for i = 1:numfiles

    % Opens current image
    filename = char(inputfilenames(i));
    image = imread(filename);
    
    % Intensity (and optionally gamma) adjust to improve contrast
    gamma = 1; % gamma value, empirical based on starting image contrast
    image_adj = imadjust(image);
    image_adj_gamma = imadjust(image_adj, [], [], gamma);

    % Find Otsu threshold and generate binary image
    bckgrnd_lvl = graythresh(image_adj_gamma);
    binaryimage = imbinarize(image_adj_gamma, bckgrnd_lvl);

    % Erode binary image to remove pixels in tubules and sheet edges
    scalefactor = 6; % scale factor, empirical based on tubule/pixel size
    se = strel('disk',scalefactor,0);
    erodedimage = imerode(binaryimage,se);

    % Performs quantifications on images generated
    image_area = numel(image_adj_gamma);
    binary_ER_area = sum(binaryimage, 'all');
    sheet_face_area = sum(erodedimage, 'all');
    tubule_and_edge_area = binary_ER_area - sheet_face_area;

    % Outputs data to cell array
    outputdata{i+1, 1} = filename;
    outputdata{i+1, 2} = image_area;
    outputdata{i+1, 3} = binary_ER_area;
    outputdata{i+1, 4} = tubule_and_edge_area;
    outputdata{i+1, 5} = sheet_face_area;
    
% %     % OPTIONAL: display images for quality checking
% %     
%     figure % original image vs. gamma adjusted
%     imshowpair(image, image_adj_gamma, 'montage');
%     
%     figure % gamma adjusted vs. thresholded binary image
%     imshowpair(image_adj_gamma, binaryimage, 'montage')
%     
%     figure % thresholded binary image vs. eroded image
%     imshowpair(binaryimage, erodedimage, 'montage');
%     
%     figure % overlay of binary image and eroded image
%     imshowpair(binaryimage, erodedimage, 'ColorChannels', 'green-magenta');
    
end

outputdata
