## Supplementary material for "A Flat Protein Complex Shapes Rough ER Membrane Sheets": Script S2

% Manders_Overlap_With_Rotation.m

% automatically Otsu thesholds images, calculates the Manders overlap, 
% and then rotates channel 1 by 90 deg clockwise, and calculates 90 deg
% rotated Manders overlap

% returns:
% image file name
% total image area
% M1 (fraction of channel 1 signal overlapping channel 2 mask)
% M2 (fraction of channel 2 signal overlapping channel 1 mask)
% M1_90rot (fraction of 90 degree rotated channel 1 signal overlapping channel 2 mask)
% M2_90rot (fraction of channel 2 signal overlapping 90 degree rotated channel 1 mask)

% requires two vertical cell arrays of strings, which specify the file 
% names of the channel 1 (c1) and channel 2 (c2) images, listed in the same
% order, such that the first entry of the list c1_inputfilenames is the
% matching image to the first entry of the list c2_inputfilenes, and so on

% Notes to the User:
% * MATLAB must have the Image Processing Toolbox installed.
% * Images must have high contrast for the segmentation to be accurate.
% * Images must be cropped to ROIs containing only resolvable (2D) ER and
%   must NOT contain regions outside the cell. (Otherwise, the rotated
%   control comparison is meaningless).
% * Comment/uncomment the final "optional" section depending on if you 
%   want MATLAB to show image processing intermediates. It is useful to 
%   spot check a few images prior to analyzing a batch to verify quality.

% Input file names
c1_inputfilenames = {'myfile1_channel1.tif'
'myfile2_channel1.tif'
'myfile3_channel1.tif'};
c2_inputfilenames = {'myfile1_channel2.tif'
'myfile2_channel2.tif'
'myfile3_channel2.tif'};

numfiles = length(c1_inputfilenames);

% Generates an output cell array
outputdata = cell(numfiles + 1, 6); % +1 to allow space for header line
outputdata{1, 1} = 'Image';
outputdata{1, 2} = 'Image Area (pixels)';
outputdata{1, 3} = 'M1';
outputdata{1, 4} = 'M2';
outputdata{1, 5} = 'M1_90rot';
outputdata{1, 6} = 'M2_90rot';

for i = 1:numfiles

    % Opens channel 1 and channel 2 files and adjusts intensities
    c1filename = char(c1_inputfilenames(i));
    c2filename = char(c2_inputfilenames(i));
    c1 = imadjust(imread(c1filename));
    c2 = imadjust(imread(c2filename));
    image_area = numel(c1);
    
    % Finds Otsu threshold and generates binary image
    c1_binary = uint16(imbinarize(c1)); % Otsu threshold as integer matrix
    c2_binary = uint16(imbinarize(c2)); % Otsu threshold as integer matrix
    
    % Above-threshold pixel values retained, below-threshold set to zero
    c1_abovethreshold = c1.*c1_binary;
    c2_abovethreshold = c2.*c2_binary;
    
    % Manders calculation:
    % M1 = fraction of signal from channel 1 overlapping the channel 2 mask
    % M2 = fraction of signal from channel 2 overlapping the channel 1 mask
    c1_abovethreshold_signal = sum(c1_abovethreshold(:));
    c2_abovethreshold_signal = sum(c2_abovethreshold(:));
    
    c1_overlapping_c2 = c1_abovethreshold.*c2_binary;
    c2_overlapping_c1 = c2_abovethreshold.*c1_binary;
    
    c1_overlapping_c2_signal = sum(c1_overlapping_c2(:));
    c2_overlapping_c1_signal = sum(c2_overlapping_c1(:));
    
    m1 = c1_overlapping_c2_signal / c1_abovethreshold_signal;
    m2 = c2_overlapping_c1_signal / c2_abovethreshold_signal;
    
    % 90 clockwise degree rotation of channel 1
    c1_abovethreshold_90deg = imrotate(c1_abovethreshold, -90);
    c1_binary_90deg = imrotate(c1_binary, -90);
    
    % Manders calculation for the rotated image set, calculated as above    
    c1_90deg_overlapping_c2 = c1_abovethreshold_90deg.*c2_binary;
    c2_90deg_overlapping_c1 = c2_abovethreshold.*c1_binary_90deg;
    
    c1_90deg_overlapping_c2_signal = sum(c1_90deg_overlapping_c2(:));
    c2_90deg_overlapping_c1_signal = sum(c2_90deg_overlapping_c1(:));
    
    m1_90 = c1_90deg_overlapping_c2_signal / c1_abovethreshold_signal;
    m2_90 = c2_90deg_overlapping_c1_signal / c2_abovethreshold_signal;

    % Outputs data to cell array
    outputdata{i+1, 1} = c1filename;
    outputdata{i+1, 2} = image_area;
    outputdata{i+1, 3} = m1;
    outputdata{i+1, 4} = m2;
    outputdata{i+1, 5} = m1_90;
    outputdata{i+1, 6} = m2_90;
    
%     % OPTIONAL: display images for quality checking
%     % Comment out when running batches!!!
%     close all
%     figure
%     imshowpair(c1, c2)
%     figure
%     imshowpair(c1_binary, c2_binary)
%     figure
%     imshowpair(c1, c1_overlapping_c2)
%     figure
%     imshowpair(c2, c2_overlapping_c1)
%     figure
%     imshowpair(c1_binary_90deg, c2_binary)

end
